# Desalination and temperature increase will shift seasonal grazing patterns of invasive *Gammarus tigrinus* on charophytes

**DOI:** 10.1101/2021.03.01.433077

**Authors:** M. Berthold, C. Porsche, A. Hofmann, P. Nowak

**Affiliations:** Applied Ecology and Phycology, University of Rostock, Rostock, Germany; Phytoplankton Ecophysiology, Mount Allison University, Sackville, Canada; Aquatic Ecology, University of Rostock, Rostock, Germany

**Keywords:** charophytes, grazing, *Gammarus tigrinus*, temperature, salinity, climate change

## Abstract

Charophytes are a refuge for zooplankton and stabilize sediments, but they are also a food source for various animal species (water birds, fishes, invertebrates). Especially the introduction of new species, like *Gammarus tigrinus*, into the Baltic Sea led to yet not understood changes in the food web. Furthermore, future projections point to increased water temperatures at lowered salinity levels affecting species capacity to acclimatize to changing abiotic factors. In this study we investigated the influence of temperature and salinity on the grazing pressure of *Gammarus tigrinus* on two charophyte species: *Chara aspera* and *Chara tomentosa*. The grazing experiments were conducted in a full factorial design with the factors salinity (3 – 13 g kg^-1^), temperature (5 – 30 °C), and charophyte species. Grazing rates were determined as mass deviation within 48 hours considering biomass changes in the presence and absence of gammarids. Grazing rate were further used to calculate charophyte losses in two coastal lagoons with different salinity concentrations for recent and future time periods. The potential grazing peak of about 24 °C is not yet reached in these coastal waters but may be reached in the near future as shown by our future projection results. However, the temperature increase, and desalination will cause a shift in seasonal individual grazing patterns from summer to spring and autumn. Desalination and temperature increase can lead to a shift in optimal habitats for *G. tigrinus* in the future.

## Introduction

Global climate change imposes several interacting effects on aquatic organisms. Besides the increase of water temperature, there may be regionally altered hydrological flow regimes that alter timing and amount of freshwater input into coastal waters. Native species get stressed by this development and ecosystems may be more susceptible to invasive species colonization. The Baltic Sea is such a system prone to eutrophication and global climate change, due to already increased sea surface temperatures, compared to other oceans worldwide (Reusch et al. 2018). The Baltic Sea is subject to large phytoplankton blooms since decades, with the riparian states working to reduce the impact of nutrient-run-off to the sea (Janssen et al. 2004, HELCOM 2018). Similar, the coastal water bodies of the Baltic Sea were impacted by eutrophication and shifts on primary producer dominance, i.e. from submerged macrophytes to phytoplankton (Schiewer 2007).

Charophytes are widely distributed in fresh and brackish water ecosystems (e.g. Schubert and Blindow 2003) and act as keystone organisms in shallow water ecosystems of the Baltic Sea because of their high biomasses (Lindner 1972). During the last decades, charophytes received attention as indicator species for oligotrophic water bodies, but also as pioneer species in post-eutrophied water bodies (e.g. Melzer 1999, Schubert and Blindow 2003).

These traits make charophytes interesting foundation species as habitats are colonized regardless of trophic status (Schubert et al. 2018). Algae of the Characeaen family are currently listed as ‘endangered species’ in several European areas (Stewart and Church 1992, Blazencic et al. 2005, Gärdenfors 2010, Auderset Joye and Schwarzer 2012) and they are also under pressure in the Baltic Sea. Comparisons of historical and recent distributions in the northern Baltic found that *C. aspera* and *C. tomentosa* were found 8 and 86% times less in the most recent transect, respectively (Pitkänen et al. 2013). Besides eutrophication related issues, salinity and/or temperature changes may influence successful recolonization in the future. Salinity changes in brackish water are such a factor, where increased salinity prevents the growth of macrophyte species (Lindner 1972). It was assumed that charophytes have several turbidity-reducing traits, like through for example lowering of resuspension of particulate matter, and direct (dissolved nutrients) and indirect (refuge for zooplankton) competition with phytoplankton (Kufel and Kufel 2002). Therefore, recolonization with those species could create feedback mechanism, favoring a macrophyte-dominated stable state.

However, restoration measures may be hampered by global climate change and the effects of desalination and increasing water temperature. Furthermore, there are hypotheses that not only abiotic parameters, but also top-down control (i.e. grazing) may affect the restoration of the pristine trophic state and macrophyte recolonization (Körner and Dugdale 2003, Östman et al. 2016). Besides grazing from native species (Körner and Dugdale 2003), there is also an increasing grazing pressure of invasive species introduced through, e.g. ballast water of ships. One of these species is the gammarid *Gammarus tigrinus* (Sexton 1939), a successful new species introduced from North America to European waters (Berezina 2007). This gammarid was described to colonize shallow waters, especially within reed, and soft bottom systems of the Baltic Sea (Daunys and Zettler 2006, Kotta et al. 2011). Amid global climate change, invasive species can alter known food web interactions and may counteract recolonization of former native primary producers (Puntila 2016). However, even these species are subject to global climate change and changes in the abiotic environment may support these invasive species even further (Kelley 2014).

The question evolves, if recolonization of macrophytes is not only hampered by abiotic factors, like temperature and salinity, but additionally by grazing pressure of (newly introduced) grazers. This grazing pressure likely changes throughout the year and may result in high-stress situations, where pressures of salinity and temperature amplify with grazing. *Gammarus tigrinus* and charophytes occur within the same depth distribution, and grazing can potentially happen. It was hypothesized that there is either a combination of factors (abiotic and biotic) that amplify mass loss on charophytes, or that some abiotic factors quench each other, e.g. lower salinity amplifies/buffers effects of higher temperatures. We tested the impact of salinity and temperature as well as their interactions on biomass change of charophytes in the presence and absence of grazers. These results also allow to discuss future developments within coastal water bodies at e.g. rising water temperatures, or desalination. Shallow coastal water bodies are likely to be more affected by such events due to their relatively shallow water column and close coupling with the adjacent land.

We chose the Darss-Zingst lagoon system (DZLS) as model ecosystem for the southern Baltic Sea, due to its salinity and trophic gradient and decade-long monitoring record (see Schiewer 2007). Even though nutrient concentrations were reduced to a comparable level between today and the 1930ies, no dense macrophyte cover was able to form in DZLS (Gessner 1957, Berthold et al. 2018a, Paar et al. 2021). Indeed, the maximum depth distribution of charophytes has not increased significantly in the DZLS during the last years, and charophytes are found only in very shallow waters, where light is not limiting (Piepho 2017). To account for this development, we combined the laboratory determined grazing potentials with field-based growth patterns of submerged macrophytes of the DZLS, and projected them with future salinity and temperature developments in these waters.

## Material and Methods

### Experimental design

We chose a full factorial design with three factors (Figure S1), for which one factor relates to species and includes two levels (*Chara aspera* and *C. tomentosa*), and the other two factors each with four levels are water temperature (9, 15, 24, and 30 °C), and salinity (3, 5, 7, 13 g kg^-1^). These salinity levels represent the salinity gradient from the Western belt Sea to the Gulf of Finland, covering most of the Baltic Sea area (HELCOM 2010). The water temperature levels represent the blooming period of shallow coastal waters (+1 °C in winter to +22 °C in summer, Schiewer 2007), with 30 °C as prospective endpoint of possible charophyte growth.

The DZLS is formed by four consecutive shallow lagoons, and shows a salinity and trophic gradient from the main river inflows Recknitz and Barthe (inner and middle part, salinity of 1 – 6 g kg^-1^, eutrophic) to the open Baltic Sea (salinity of 10 – 12 g kg^-1^, mesotrophic, Schumann et al. 2006). Salinity varies from 4 – 6 g kg^-1^ at the respective sampling locations in the Bodstedter Bodden (54°22’30,5" N, 12°34’14,9" E) and around 6 – 8 g kg^-1^ in the Grabow (54°22’2,3" N, 12°48’27" E) (Schumann et al. 2006). During the experiment, two charophyte species were used, *Chara aspera* and *Chara tomentosa*. Both species are common in the Baltic Sea, with *C. aspera* covering the widest salinity range among charophytes (Blindow 2000). Along the west coast of Sweden, *C. aspera* is regularly found at sites with salinities up to 15 g kg^-1^ (temporarily up to 20 g kg^-1^), but it is also common in freshwater (Hasslow 1931, Blindow 2000). *C. tomentosa* is among the largest charophyte species and can be found in freshwaters all over the world. Only in the Baltic Sea, *C. tomentosa* also occurs in brackish water, with a salinity range between 0 – 7.5 g kg^-1^ (Björkman 1947, Torn et al. 2003). Gammarids were derived from the Aquaculture section in Born of the State Research Centre for Agriculture and Fishery Mecklenburg-Vorpommern. This research facility is located at the DZLS and uses brackish water from the lagoon with a salinity of 4 – 6 g kg^-1^. Gammarids were identified by morphology and genetic analyses (see supplement).

Charophytes were extracted as intact plants from the DZLS. Charophytes were then cultured in beakers (vol. 500 ml) following the culture design described by Wüstenberg et al. (2011). Each beaker contained sediment containers (vol. 64 ml) with acid-rinsed, phosphorus-enriched sediment (6 g P kg^-1^ sediment). Mineral growth media was prepared to culture charophytes (Pohl et al. 1987, Wüstenberg et al. 2011), but was adjusted to the respective salinities by adding marine salt (Instant Ocean, Aquarium Systems, Sarrebourg). Salinity was checked with a salinometer (Multiline P4, WTW). Each of the 16 combinations were replicated eight times with gammarids (= grazing treatment) and in absence of gammarids (= control) (Fig. S1). Gammarids were pre-acclimatized to experimental conditions by culturing them at the respective temperature and salinity combinations. Ten to 20 individuals were simultaneously cultured per temperature-salinity combination, aerating all cultures.

Charophytes were added as food supplement, as well as resting surface for the gammarids.

Charophytes were wet weighed and set into beakers filled with sediment and growth media of the respective salinity. Controls consisted of one charophyte, whereas grazing treatments consisted of one charophyte and five gammarids in one beaker. Control and grazing treatments were exposed to light intensities between 80 and 120 μmol photons m s using fluorescent tubes (Phillips, TLD 36W/950). Each replicate run lasted for about 48 hours.

Replicates were run in blocks over a period of four weeks per *Chara* species. Two out of the eight replicates per salinity-temperature combinations were conducted simultaneously to spread and randomize replicate runs over the four week period. According to previous studies on growth of charophyte (Nowak et al. 2017) we did not expect significant growth responses of *C. aspera* and *C. tomentosa* over the course of two days. However, it was expected to find significant mass deviations when gammarids were present.

Charophytes were wet weighed after each replicate run to calculate mass differences. Dead gammarids were counted at the end of each replicate run and replaced for the next replicate run. If possible, the same gammarids were used throughout all replicate runs. Equations 1 – 3 were used to extract grazing rates of gammarids from growth rates of charophytes during the experiments. Grazing rates were treated with an error-propagation to account for experimental variability.

Equation 1 subtract the mean effect of biomass change of the control (C) from the treatment which also includes grazing (T):

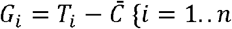

Equation 2 standardizes the grazing effect from prior formula (**z**-score):

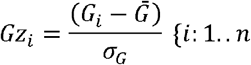

Equation 3 calculates the mean of pure grazing and include its error caused by the experiment:

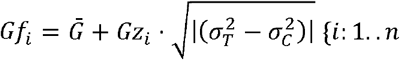

T = treatment, C = control, G= detrended grazing, G_z_ = standardized grazing, G_f_ = grazing with included experimental error, n = replicates

### Field sampling and analyses

In addition, we used elemental and biomass composition of submerged macrophytes from the DZLS, to compare the experimental derived grazing rates to the actual ecosystem. Field sampling was conducted in the Bodstedter Bodden and the Grabow in 2014, as part of the BACOSA project. Interestingly, the Bodstedter Bodden has on average a lower salinity of 2 g kg^-1^, then the Grabow, coinciding with future desalination projections (Neumann and Friedland 2011). This difference may be used as model validation, if grazing rates of the present Bodstedter Bodden represent future grazing rates in the Grabow. Submerged macrophytes were sampled throughout the year, first in 54x54 cm plots, and in 25×25 cm plots later during the year at nine different depths. Here, three sampling locations were pooled into three depth zones, according to water depth and proximity to the reed belt. Water depths for both stations varied with water levels in the respective lagoons and were shallow (30 – 60 cm), intermediate (60 – 90 cm), and deep (90 – 120 cm). Sampled macrophytes were separated on the genus level. Whole plants, including roots, were wet-weighed and then dried at 60 °C for 24 h. Afterwards, dry mass (DM) was re-weighed again to calculate water content. Dried plant material was ground with a ball mill. The powder was treated with 1M HCl to remove inorganic carbon residues and then weighed in tin-container (1 × 0.5 cm). These samples were analyzed for their carbon and nitrogen content using an elemental analyzer (varioEL III). The analyzer was calibrated using acetanilide (∼5 mg per sample). Aliquots of dried plant material were weighed in crucibles and combusted at 550 °C for 4 h. The ash was re-weighed to calculate ash-free dry mass. Ashes were used to determine total phosphorus content (mg P g^-1^ dry mass), using an acid persulfate extraction (see Berthold et al. 2015).

### Data analyses

We further used the experimental grazing data to calculate grazing rates based on salinity and water temperature at two sites of the DZLS. The data set failed to show homogeneity of the residuals, and was therefore Box-Cox power transformed (*bcnPower* function, R package “*car*”, Fox and Weisberg 2019). Finally, general linear models (GLM, *glm* function, R package “stats”, R Core Team 2019) and generalized additive models (GAM, *gam* function, R package *“mcgv”,* Wood 2011), both with Gaussian distributional assumptions, were compared by the Akaike Information Criterion (AIC, R package “stats”, R Core Team 2019) and the Bayesian Information Criterion (BIC, R package *“stats”,* R Core Team 2019), choosing the model with low AIC and BIC values. We chose the GAM (with knots, k = 16) as the best model to reflect the experimental data (see Results). To our knowledge, recent studies on seasonal population dynamics of *G. tigrinus* are currently missing in our study system, as elsewhere in the Baltic Sea. However, recent field studies found 5 to 2200 Ind m^-2^ (gammarids on genus level) in the DZLS and the adjacent Western Rügen lagoon system.

Largest, but fewest animals were found early in the year (22 – 28 mg Ind^-1^), with decreasing mass, but increasing abundance in summer (3.5 mg Ind^-1^) (M. Paar, personal communication). These patterns point to the same seasonal development as described in Chambers (1977). If we assume that *G. tigrinus* constitutes 20% of the local gammarid population in the DZLS (Zettler 2001), we get a population size of up to 450 Ind m^-2^. These numbers are in agreement with first described abundances of *G. tigrinus* of 300 – 500 Ind m^-2^ in the phytal and reed belt of the DZLS (Zettler 1995), or 30 – 110 Ind m^-2^ in the Neva estuary (Baltic Sea, Berezina 2007).

We used again a GAM model to calculate the seasonal population development of *Gammarus tigrinus*, normalized the predicted seasonal data to have a maximum of 1. Then, we used this normalized prediction to calculate population densities with a maximum of 400 Ind m^-^² (Zettler 1995, M. Paar personal communication, Fig. S6). These population numbers were combined with the individual grazing rates of the temperature-salinity model with field data from the two sites of the DZLS (Equation 4) to calculate seasonal grazing rates. We used a fixed grazing preference of 20% macrophyte share in gammarid diets, that means grazing rates were multiplied by a factor of 0.2 (Pellan et al. 2016). Furthermore, we used the GAM to predict the charophyte development as function of sampling date (k = 8) at two sites of the DZLS. We calculated daily biomass increase and subsequently added gammarid grazing rates to determine the potential gross biomass production per site (see Results and Discussion below).

Equation 4 calculates the seasonal grazing rates per square meter and day for specific sites:

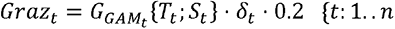

biomass change of the control t = day of the year, Graz_t_ = grazing at day t (mg·m^-^²·d^-1^), G_GAMt_ = modelled grazing rate of one individual at time t with given temperature (T) and salinity (S) from specific site at time t, δ_t_ = population density at time t (individuals·m^-^²), 0.2 = reflects the 20 % fixed grazing preference, n = length of the year in days

## Results

### Growth and survival rate

Starting biomasses of *C. aspera* in control and treatment beakers ranged from 50 to up to 300 mg, with the most macrophytes being within the range of 100 – 175 mg (70 %). After two days, biomass distribution of *C. aspera* in control beakers stayed relatively the same, but smaller and larger biomass proportions increased. Biomass of *C. aspera* decreased with increasing temperature at all salinities, but strongest at salinities > 7 g kg^-1^. Contrary, biomass stayed almost constant at lower temperature, even at higher salinities (Supplement Table S2 and Figure S4).

Starting biomass of *C. tomentosa* in control beakers ranged from 75 to up to 600 mg, and 40 % of all macrophytes were within the range of 100 – 125 mg at the beginning. Contrary to *C. aspera*, biomass of *C. tomentosa* in treatment beakers was close to a normal distribution with 50 % of all macrophytes being within the range 175 – 225 mg at the beginning. The biomass distribution of *C. tomentosa* in control beakers stayed almost the same, but with a trend to smaller biomasses, indicating minor biomass losses even without grazing. Biomass of *C. tomentosa* stayed very constant at almost every temperature-salinity combination. There was a possible growth at temperatures of 15 to 24 °C at salinities of 7 g kg^-1^ (Supplement Figure S4). If results of both charophyte control beakers were combined, the area with stable biomass over two days expands from 10 – 25 °C and salinities of 7 – 13 g kg^-1^. In total, these results indicate an overall stable experimental approach, as significant growth was not expected within two days.

Two thirds of the time (65 %) all gammarids survived at all temperature-salinity-macrophyte combinations in treatment beakers. Additionally, 30 % of the time, only one gammarid died during incubation. These results indicate overall stable culture conditions and the capability of these gammarids to survive a wide range of temperature-salinity gradients.

### Gammarid grazing on charophytes

Biomass in treatment beakers decreased in all combinations and beakers. Differences to control beakers were always significant, regardless of charophyte species, or treatment (see Supplement table S2). Grazing rates ranged between 1 to 10 mg FM Ind^-1^ gammarid d^-1^ on *C. aspera*, already accounting for biomass changes in control beakers. Grazing rates peaked at 24 °C and tended to be higher with increasing salinities. Grazing ranged between 1 to 5 mg FM Ind^-1^ gammarid d^-1^ on *C. tomentosa*. Grazing rates peaked again at 24 °C for all salinity levels but with a tendency to slightly lower salinity compared to grazing rates on *C. aspera*. Interestingly, highest grazing rates per charophyte species fell into temperature ranges where charophytes already showed highest loss rates in control beakers. In general, grazing rates on *C. aspera* were up to two times higher compared to *C. tomentosa*. Pooled grazing rates of both charophyte species were on average up to 5.5 mg Ind^-1^ d^-1^.

### Field data results and model

Seasonal daily grazing rates within the ecosystem would range from 0.1 up to 5 mg FM Ind^-1^ d^-1^, when applying the grazing model on field observations of temperature and salinity of Grabow and Bodstedter Bodden (Figure 5). Grazing rates would peak during summer and be lowest at winter at the two sampling stations of the DZLS (see Figure 5). *Gammarus tigrinus* was originally collected in April/May, where grazing rates are not at its peak. Nonetheless, gammarids grazed twice as much at 24 °C in our treatments, as would be expected from their initial collection time in April and May. These findings indicate acclimation processes.

Interestingly, the modelled grazing rates differed significantly between both stations (p < 0.001, paired Mann-Whitney U-Test), even though the difference in salinity are only 2 g kg^-1^. Charophyte biomass was highest in the shallow areas, close to the reed belt at both sampling sites (Figure 6). Biomass in the shallow zone peaked in July in the less eutrophic Grabow, and in October in the highly eutrophic Bodstedter Bodden. Contrary, charophyte biomass was higher in the Bodstedter Bodden at intermediate distances/water depths compared to the Grabow. However, Grabow biomass peaked during summer, whereas biomass in the Bodstedter Bodden became depressed during that time. Charophytes were only found in very little biomasses in the deeper water areas at both sampling locations, indicating a possible light limitation from shallow to deep.

Charophyte biomass would have been up to 15 – 50% higher at shallow and intermediate depths, if macrophyte growth is corrected for seasonal gammarid grazing. Additionally, gammarid grazing would follow charophyte biomass peaks at the respective stations. The yearly potential grazing rate for *G. tigrinus* on charophytes ranges from 37.2 to 42.5 g FM m^- 2^, considering regional salinity and temperature patterns within the DZLS.

### Future projections on desalination and temperature increase in the period of 2050 – 2100

The projected grazing rates vary significantly from today’s grazing rates (p < 0.001, paired Mann-Whitney U-Test), when considering future desalination and temperature increase in this region (van der Linden and Mitchell n.d., Neumann and Friedland 2011). Individual grazing rates would increase in spring and autumn at both stations, but grazing rates in the Grabow being at least twice as high as in the Bodstedter Bodden. Furthermore, individual grazing rates would probably drop during summer, as water temperatures will fall out of optimum ranges for *G. tigrinus*. These differences in future grazing rates are caused by earlier warmer temperatures, and lower salinities, increasing future individual grazing pressure in the Grabow. However, this double peak is not apparent, when the seasonal population structure is included. Areal grazing rates would not be different during spring, and even lower during summer. However, grazing rates would strongly increase during autumn at both stations. This peak is caused by the late population peak of gammarids, and the increased temperature in future projections. This grazing pattern would likely affect charophytes in the turbid Bodstedter Bodden, as they showed highest biomass during autumn.

## Discussion

Charophyte growth and gammarid grazing rates in our experiment and in the field depend on changes in the abiotic environment. However, grazing rates are simultaneously influenced by available food sources for gammarids, occurrence of competing species and predators, or ultimately habitat choice and niche occupation.

In this study, desalination and temperature increase experiments lasted only for two days. Charophytes of control beakers showed only little, if any biomass increase, and cannot be used for long-term growth projections. There are complex interactions between abiotic factors (e.g. salinity, temperature, light climate) that influence charophyte growth and these effects depend on species-specific characteristics, local environmental conditions, and probably locally adapted sub-populations (Blindow et al. 2009, Auderset Joye and Rey-Boissezon 2015, Rojo et al. 2017). Distinct *C. aspera* populations can show different growth optima depending on the habitat (freshwater and brackish), which is translated by changing photophysiological characteristics at varying salinities (Blindow et al. 2003, Blindow and Schütte 2007). Likewise, water temperature shapes charophyte populations, which may result in local biomass decreases of *C. aspera*, *C. tomentosa* and *C. vulgaris* if temperature increases more than 2 °C (Auderset Joye and Rey-Boissezon 2015, Choudhury et al. 2019). Even our short-term experiments may confirm these results, as both charophyte species lost most biomass at temperatures above average summer habitat temperatures (> 24 °C). However, interactions of abiotic factors do not necessarily induce synergistic effects in charophytes (Rojo et al. 2017, Puche et al. 2018). Our short-term growth controls showed synergistic effects on charophyte growth rates for different temperature and salinity conditions (Fig S4). Further studies are needed to clarify long term charophyte changes in the face of increasing water temperatures and decreasing salinities, especially among locally adapted sub- populations.

Several physiological reactions, like survivability or oxygen consumption of *G. tigrinus* on salinity and temperature changes have already been described in the literature (Dorgelo 1973, 1974, Koop and Grieshaber 2000). *Gammarus tigrinus* survives within a broad salinity-temperature spectrum, where only full marine or freshwater conditions show highest mortalities. Furthermore, potential synergetic were observed with higher temperatures buffering higher salinities, with survival rates 10-times lower at 5 °C, compared to 25 °C (Dorgelo 1974).

Grazing rates in our study suggest similar findings, as lowest temperatures strongly reduced grazing rates, regardless of salinity. There was also a tendency to higher grazing rates, at higher salinity/temperature combinations, compared to high salinity/low temperatures, indicating a grazing optima at higher temperatures. Ion regulation in *G. tigrinus* happens fast, and can regulate sudden high ionic stress to a variety of ions (Na, Cl, K) without compensation losses. However, hypoosmotic, that means low salinity conditions, cause increased NO_3_ inflow into *G. tigrinus* hemolymph, probably causing decreased survivability (Koop and Grieshaber 2000). The growth media for charophytes in our experiment has elevated NO_3_ concentrations (3 mM KNO_3_, Pohl et al. 1987). This elevated NO_3_ concentrations may have caused lower grazing rates at lower salinities in our study.

Besides abiotic factors, population structure influenced the grazing rates of gammarids. Populations of *G. tigrinus* were described to turn-over at least three times per year, with a peak of large animals (6 – 9 mm) in spring, and a concomitant shift to smaller animals in late summer (2 – 4 mm) (Chambers 1977). We used animals in the size range of 6 – 12 mm, i.e. large adults for all of our experiments. This size class is only dominant by up to 60% until May, and is almost completely replaced by smaller animals from June to September (Chambers 1977). This pre-selection had likely an effect on the described grazing rates, as metabolic rates of *G. tigrinus* increase with lower weight in form of a power function (Normant et al. 2007). Gammarids in our study weighed from 10 – 90 mg wet mass Ind^-1^ (data not shown), whereas the gammarids of Normant et al. (2007) were caught in August, and weighed only 5 – 25 mg wet weight Ind^-1^. Animals of our study should therefore showed lower specific metabolic rates of 0.25 – 1.5 mW g^-1^ wet mass, compared to 1 – 3.5 mW g^-1^ wet mass in smaller gammarids (Normant et al. 2007). This reduced metabolic rate had likely an impact on the grazing rates we found in this study, as grazing for metabolic upkeep is lower in large adult animals.

The results of this study indicate that the individual grazing rates of *G. tigrinus* will change over the next 50 to 100 years. During that time, the optimal properties of temperature and salinity for *G. tigrinus* will increase in spring and autumn months and decrease during summer months.

Springtime is a crucial period for charophyte sprouting, as the active growth of for example *C. tomentosa* takes place during a relatively short period at the beginning of summer (Torn et al. 2006). Furthermore, young sprouted charophytes show a higher elemental content per biomass, than for example tracheophytes (Volkmann 2016, this study), making them a preferred grazing opportunity (Kraufvelin et al. 2006). *Gammarus locusta* follows a preference for high value plant material and selectively feeds on more nutrient-rich macrophytes, especially periphyton, brown and green algae (Kraufvelin et al. 2006). If *G. tigrinus* shows similar grazing preferences, periphyton from tracheophytes would be the first choice followed by charophytes in the DZLS. Periphyton (Sanudo-Wilhelmy et al. 2004) and possibly charophytes (Kufel and Kufel 2002) can take up large amounts of dissolved nutrient that occur frequently through diffuse run-off (Berthold et al. 2018b). Tracheophytes in the DZLS and adjacent lagoons can show high colonization rates by epiphytes (Paar et al. 2021), offering an additional plant food source to gammarids. Nonetheless, charophytes in the DZSL showed higher C:N and N:P ratios, then tracheophytes, making them more likely to gammarid grazing as high-value food (see Fig S7). Furthermore, this pre-selection of high-value food may explain the differences in grazing rates on *C. tomentosa* vs. *C. aspera* found in this study. *C. tomentosa* can show higher C:N ratios (44:1) than *C. aspera* (27:1), making it maybe a less favorable food source. Furthermore, *C. aspera* showed in our study signs of early decay at higher temperatures, contrary to *C. tomentosa* (see trends in control beakers, Figure S4). Such beginning decay can weaken *C. aspera* and make it more susceptible to grazing (Buchsbaum et al. 1991), explaining the deviation of grazing rates between both charophytes along the temperature gradient found in this study.

The drop of grazing rates during future summer conditions is probably caused by below- optimum water temperatures in combination with lowered salinities. The impact on future summer impact grazing is more pronounced in Bodstedter Bodden, then in the Grabow. This difference is probably caused by even lower salinity concentrations in the Bodstedter Bodden, indicating that future desalination can lead to shifts of preferred habitats.

Food preferences can change throughout the year depending on gammarid life stage and temperature (Felten et al. 2008, Pellan et al. 2016). Plant biomass becomes increasingly important, when temperatures increase and can reach up to 20 – 30% of total food consumed (Pellan et al. 2016). Grazing on charophytes can happen earlier in the year, and lasts longer during autumn. This effect does not come into effect in our future projection model, as the current modelled population densities of gammarids are lowest in spring. Nonetheless, prolonged grazing may hamper charophytes sprouting in spring (*C. aspera*) and lower biomass of overwintering species (*C. tomentosa*) (Torn et al. 2006, Blindow et al. 2016).

Functional feeding in gammarids relies furthermore on population composition and food availability (Kelly et al. 2002). Other food sources like detritus, plant litter, or animal matter, like chironomides, are usually important food sources during winter time (Felten et al. 2008, Pellan et al. 2016). Sediment organic content can be as high as 20% in sediments of the DZLS (Berthold et al. 2018c). Sediment detritus was not an alternative source in our experiments, as we used only inorganic sand. Plant litter occurs abundantly within the reed belt during winter time in the DZLS (Karstens et al. 2016). Our experimental results show therefore only a potential grazing rate, as we did not represent the variety and abundance of alternative food sources from within the ecosystem.

Furthermore, grazing rates depend on niche occupation and competition with native and other invasive species. *Gammarus tigrinus* can probably outcompete the native *G. salinus*, if both depend on *Pylaiella littoralis* in the northern Baltic Sea (Orav-Kotta et al. 2009). Both gammarids had the potential to exceed daily macrophyte production during summer (Orav-Kotta et al. 2009). *G. tigrinus* may be more tolerant against heat waves and sub-oxic conditions, than native species (Lenz et al. 2011). These broader temperature acclimation abilities may help to permanently introduce *G. tigrinus* within the Baltic Sea, as sea surface temperature are already higher there, then in other ocean parts (Reusch et al. 2018). Furthermore, endurance of sub-oxic conditions is an advantage in eutrophic coastal waters of the Baltic, as redox conditions within the reed belt can change fast (Karstens et al. 2015), and an increase of nutrients is challenging submerged macrophytes in the coastal water bodies of the southern Baltic Sea (Paar et al. 2021), stressing habitats of native gammarids. Contrary, an *in silico* study assumed that *G. tigrinus* has actually a narrower niche than native gammarids in the northern Baltic Sea (Herkül et al. 2016). These results are somewhat contradictory to other studies in this area (Korpinen and Westerbom 2009, Kotta et al. 2011, Lenz et al. 2011), and may have not considered the impact of haplotype-diversity (Baltazar-Soares et al. 2017), that means locally adapted populations of *G. tigrinus*. Different studies revealed high genetic variability of *G. tigrinus* in the Baltic Sea (Supplement Table 1, Figure S2, Kelly et al. 2006 and Paiva et al. 2018) indicated that the invasive species may have different tolerance and different limits to salinities due spatially varying selection among populations.

The patchy macrophyte stands found in the DZLS, and in other eutrophic water bodies, may affect the occurrence of *G. tigrinus*, and therefore the grazing rates, especially in intermediate and deeper waters. Palaemonid prawns put a higher grazing pressure on *G. tigrinus* in unvegetated mesocosms, then in vegetated ones (Kuprijanov et al. 2015). It is likely that *G. tigrinus* stays close to the reed belt, as described for this lagoon and other coastal waters (Zettler 1995, 2001). This spatial distribution would further hamper charophyte recolonization, as the shallow areas in the DZLS are the only spots, where light availability is high enough to support charophyte biomass (see Figure S6, Piepho 2017).

## Acknowledgements

The authors thank the State Agency for Environment, Nature Conservation and Geology Mecklenburg-Vorpommern for providing monitoring data and the Aquaculture section in Born of the State Research Centre for Agriculture and Fishery Mecklenburg-Vorpommern for providing gammarids. We also want to thank Dr. Ralf Bastrop for providing the respective primers, and Dr. Martin Paar for providing us with recent data on gammarid distribution in the southern Baltic Sea.

---

**Figure 1:**
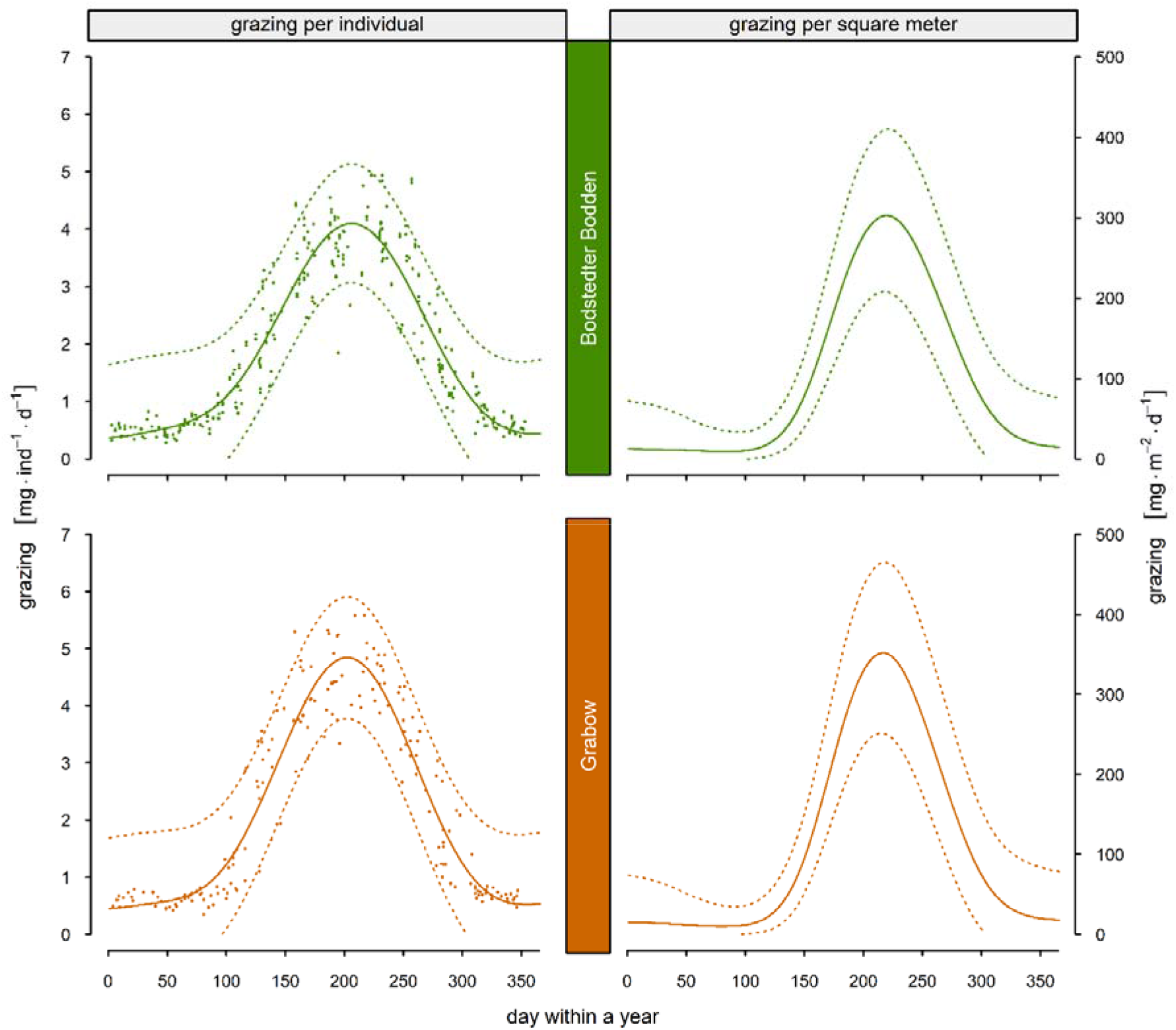
Seasonal losses of *Chara* sp. biomass by grazing of *Gammarus tigrinus* in the Bodstedter Bodden (upper panel) and Grabow (lower panel) over the years of 2000 to 2018. The losses of *Chara* sp. biomass by individual grazing (points) were modelled (generalized additive model) depending on temperature and salinity conditions at the respective time point in the water body (left). Grazing per square meter (right) was calculated as the product of the individual grazing model (left) and the seasonal population development of *Gammarus tigrinus* (Supplement Figure S5) with a maximum abundance of 400 individuals per square meter. The solid line represents the mean and the dotted line the 95 % confidence interval.

**Figure 2.**
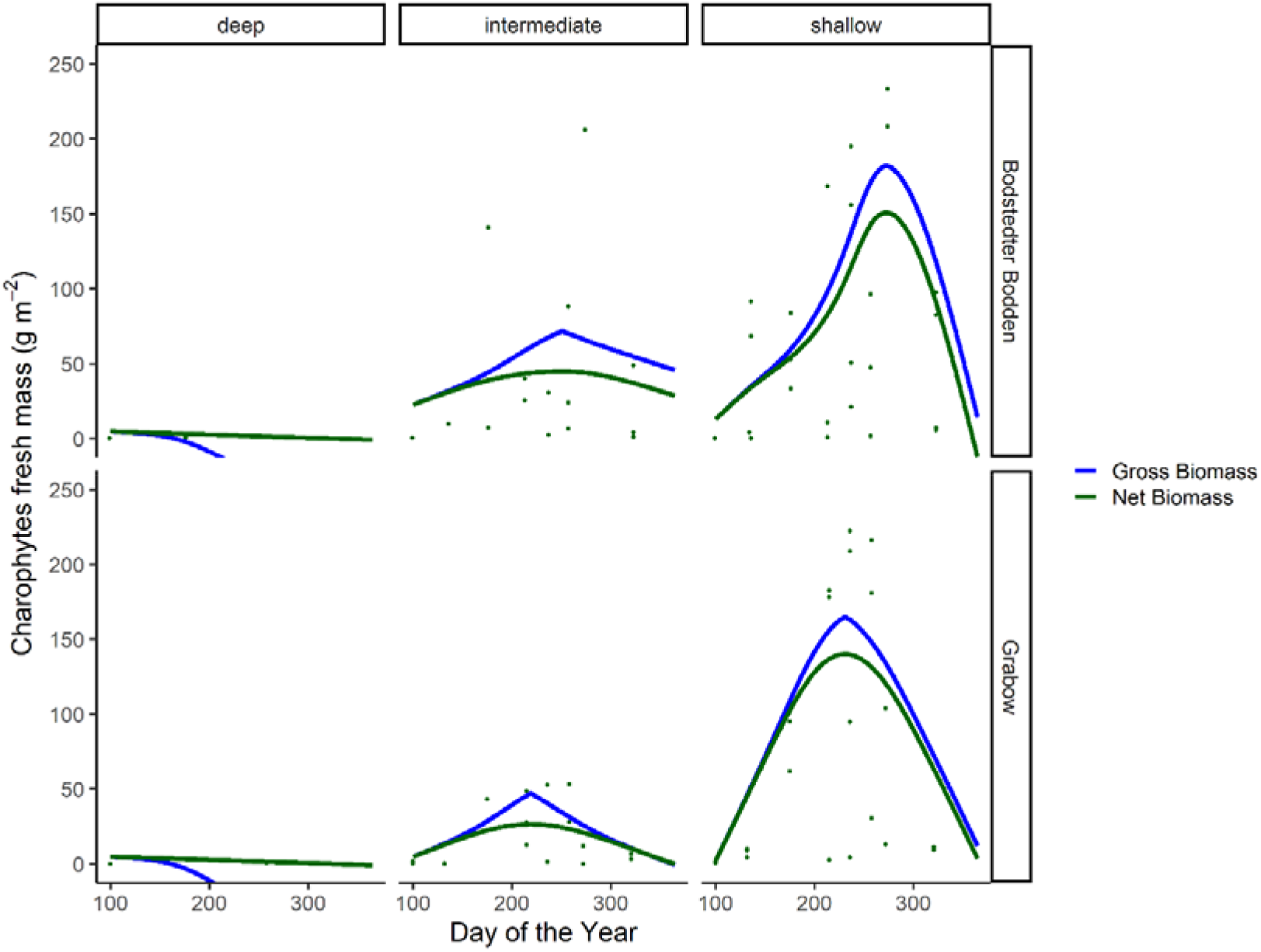
Charophyte fresh mass in gram per square meter at two sampling locations of the Darss-Zingst lagoon system along a water depth/distance to reed belt transect. Shallow depth (30 – 60 cm) and close to reed belt, intermediate depth (60 – 90 cm) and distance to reed belt, deep depth (90 – 120 cm) and farthest distance to reed belt. Net biomass represents model- fitted values (generalized additive model) of observed charophyte biomass. Green points represent observed charophyte biomass at the respective day. Gross biomass represents the sum of net biomass plus biomass possibly lost by abiotic influenced seasonally resolved *Gammarus tigrinus* grazing.

**Figure 3:**
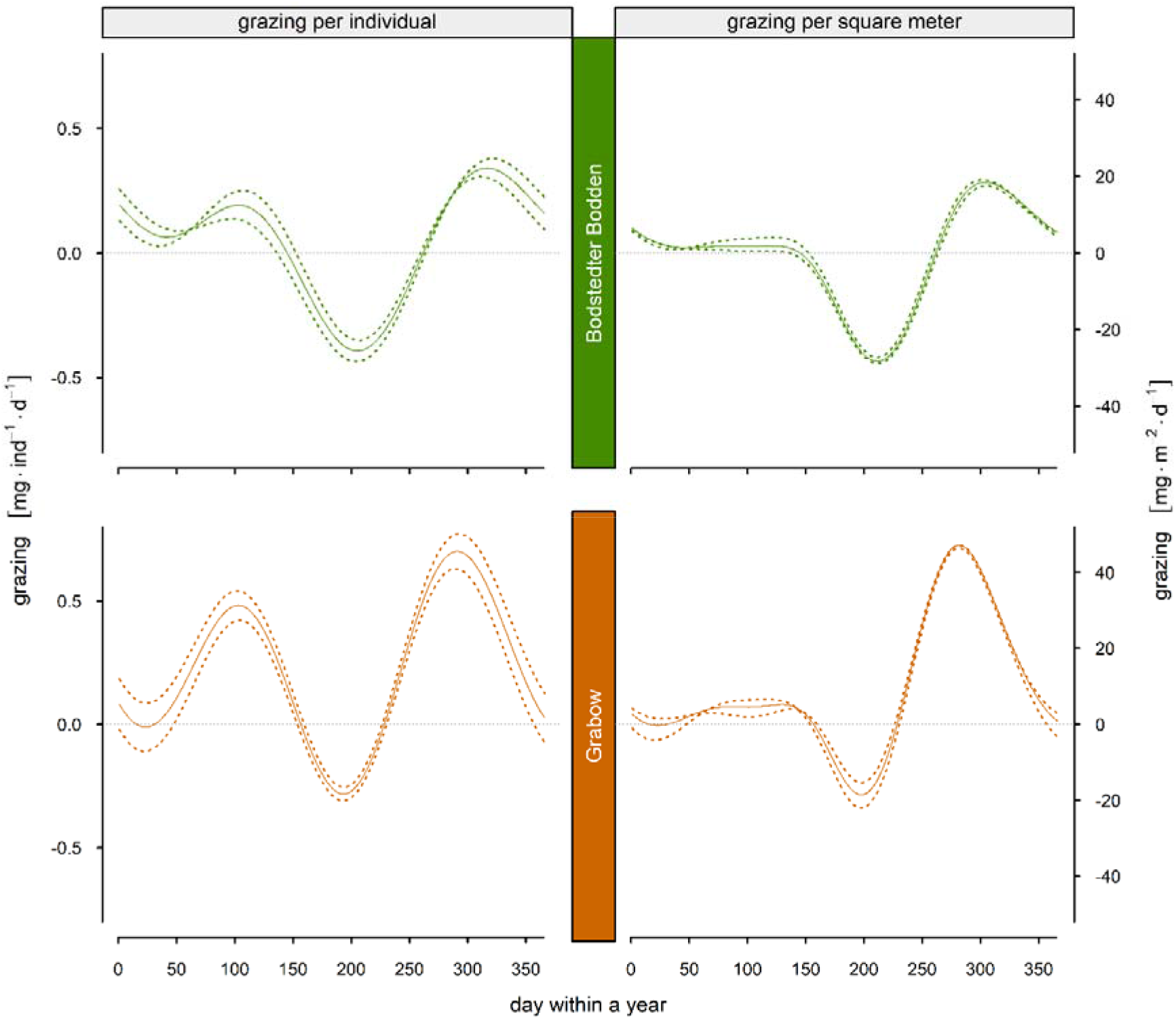
Future projection of seasonal losses of *Chara* sp. biomass by grazing of *Gammarus tigrinus* in the Bodstedter Bodden (upper panel) and Grabow (lower panel) as difference to present day grazing. Projections were calculated with a delta change approach of recent data with projected changes. Data from 2000 to 2018 was modified in temperature (+2.75 K) and salinity (-1.75 g kg^-1^) as described in Neumann and Freidland (2011) and van der Linden, P. and Mitchell, J. F. B. (2009). Losses of *Chara* sp. biomass by individual grazing were modelled depending on the modified temperature and salinity conditions at the respective time point in the water body (left). Grazing per square meter (right) was calculated as the product of the individual grazing model (left) and the seasonal population development of *Gammarus tigrinus* (Supplement Figure S5) with a maximum abundance of 400 individuals per square meter. The solid line represents the mean and the dotted line the 95 % confidence interval.

**Table 1:**
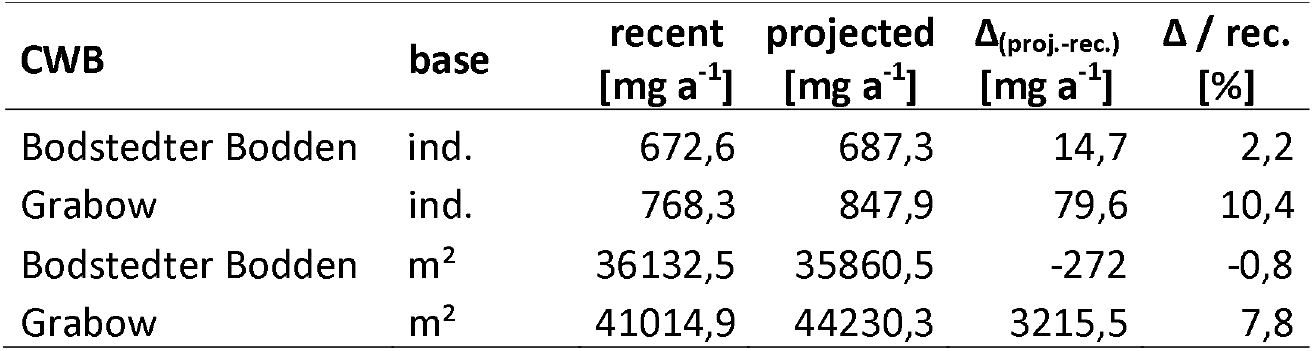
Sum of losses of *Chara* sp. biomass by individual grazing or by grazing at one square meter with respect to the seasonal population pattern of *Gammarus tigrinus* (Supplement Figure S5) with a maximum abundance of 400 individuals per square meter. These were calculated for the Bodstedter Bodden and the Grabow as well as for the recent data and the future projection.

## Supplement

### Genetic analyses

At the end of the grazing experiment, 69 gammarids were determined as *Gammarus tigrinus* following Zettler and Zettler (2017) and preserved in 80 % ethanol. To verify the morphological analyses, a genetic approach was conducted on 15 individuals. Total genomic DNA was extracted using a silica spin column procedure with the DNeasy Blood and Tissue Kit (Qiagen, Hilden, Germany) following the protocol provided by the manufacturer. Partial sequences of the COI gene were amplified with the universal primers LCO 1490 and HCO 2198 (Folmer et al. 1994).

PCR amplifications were performed with a denaturation step for 60 s at 94°C, followed by 5 cycles of: 60 s at 94°C, 90 s at 45°C and 60 s at 72°C, 35 cycles of: 60 s at 94°C, 90 s at 51°C and 60 s at 72°C, and completed with 5 min at 72°C as a final extension step. PCR was performed in a 30 µl reaction volume with a Taq PCR Master Mix (Qiagen, Hilden, Germany) consisting 2.5 mM MgCl2, and 0.5 pmol of each primer (final concentration). The PCR products were extracted from agarose gels according to the protocol of the Biometra- innuPrep Gel Extraction Kit (Analytik Jena, Jena, Germany), and were sequenced directly using a 3130×L Genetic Analyzer (Applied Biosystems, NY, USA) with sequencing primers identical to the primers that were used for the PCR reaction. Obtained sequences were quality controlled and aligned via the BIOEDIT software (Hall 1999). The COI gene is widely used for inter- and intraspecific diversification questions concerning amphipod taxa. For the taxonomic determination addition GenBank sequences of *Gammarus tigrinus* (see Supplement Table S1), *G. duebeni* Liljeborg, 1852 (EU421779), *G. locusta* Linnaeus, 1758 (KT209211), *G. oceanicus* Segerstråle, 1947 (GQ341809), *G. pulex* Linnaeus, 1758 (MN400977), *G. roeseli* Gervais, 1835 (EF570337) *G. salinus* Spooner, 1947 (KT208533), and *G. zaddachi* Sexton, 1912 (KU845083) were used for the phylogenetic analyses.

In order to estimate the evolutionary divergence between haplotypes, pair-wise uncorrected p- distances and the number of substitutions were conducted using MEGA version X (Kumar et al. 2018). To uncover phylogenetic relationships, a Maximum likelihood tree was constructed using MEGA version X (Kumar et al. 2018) on the basis of the Kimura 2-parameter model. Branch support for the nodes was calculated from 1000 bootstrap replicates.

To study the distribution of mtDNA sequence diversity, a haplotype network was constructed using the Median-joining algorithm (Bandelt et al. 1999) implemented in PopART (Leigh and Bryant 2015). The distribution basemap was created with QGIS.org software (http://www.qgis.org) and modified in CorelDRAW (Corel Corporation, Ottawa, Ontario, Canada).

**Supplement Table 1:**
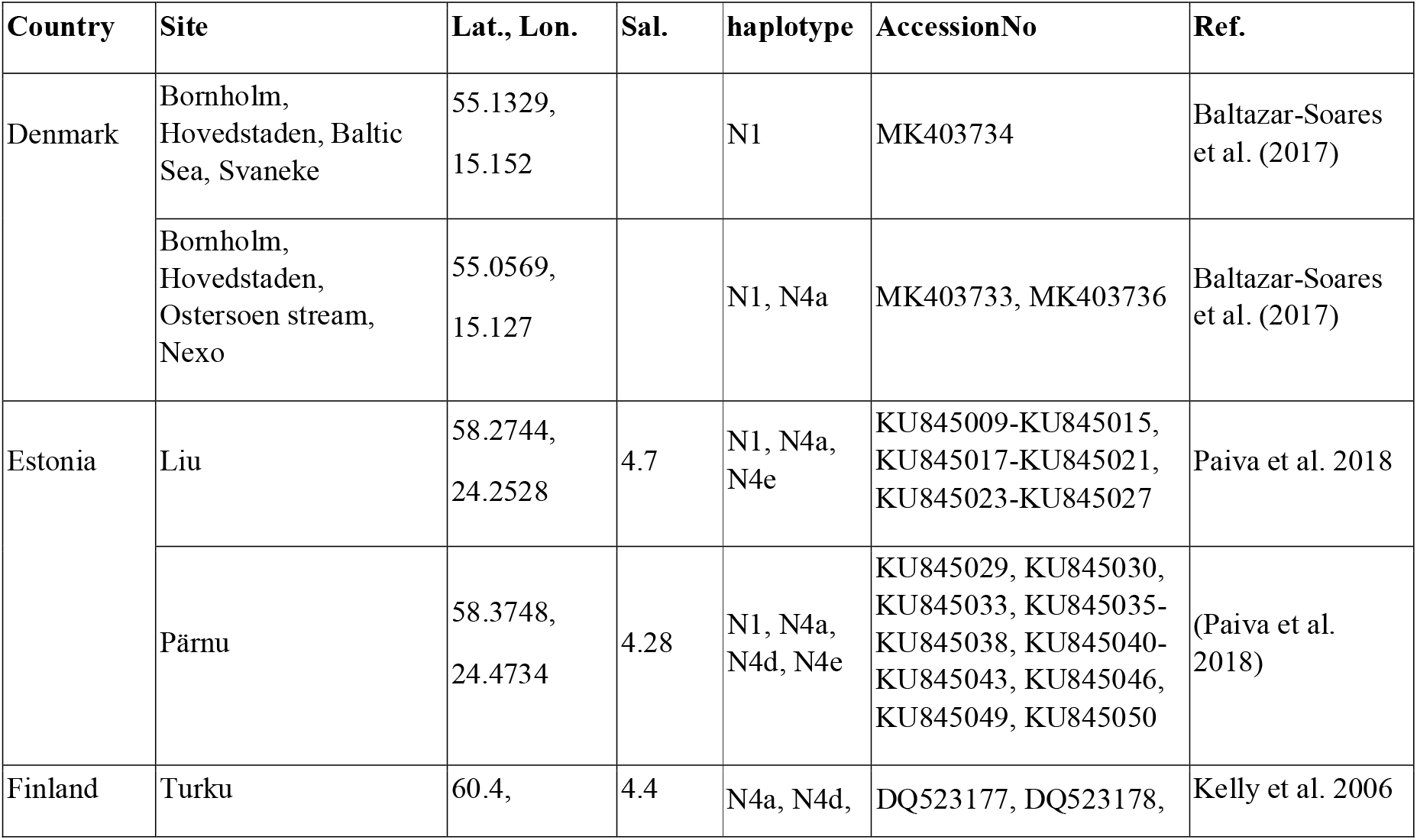
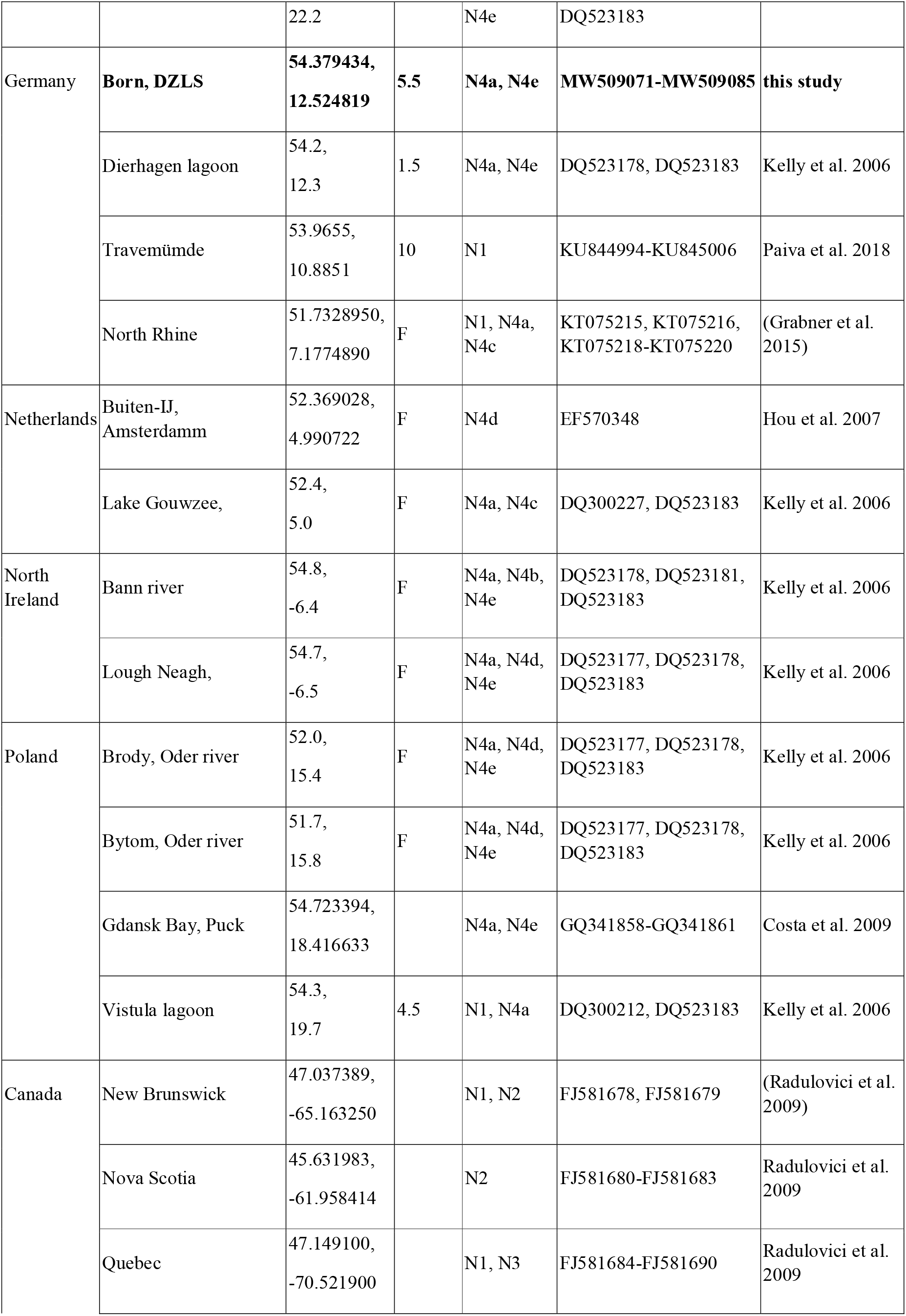
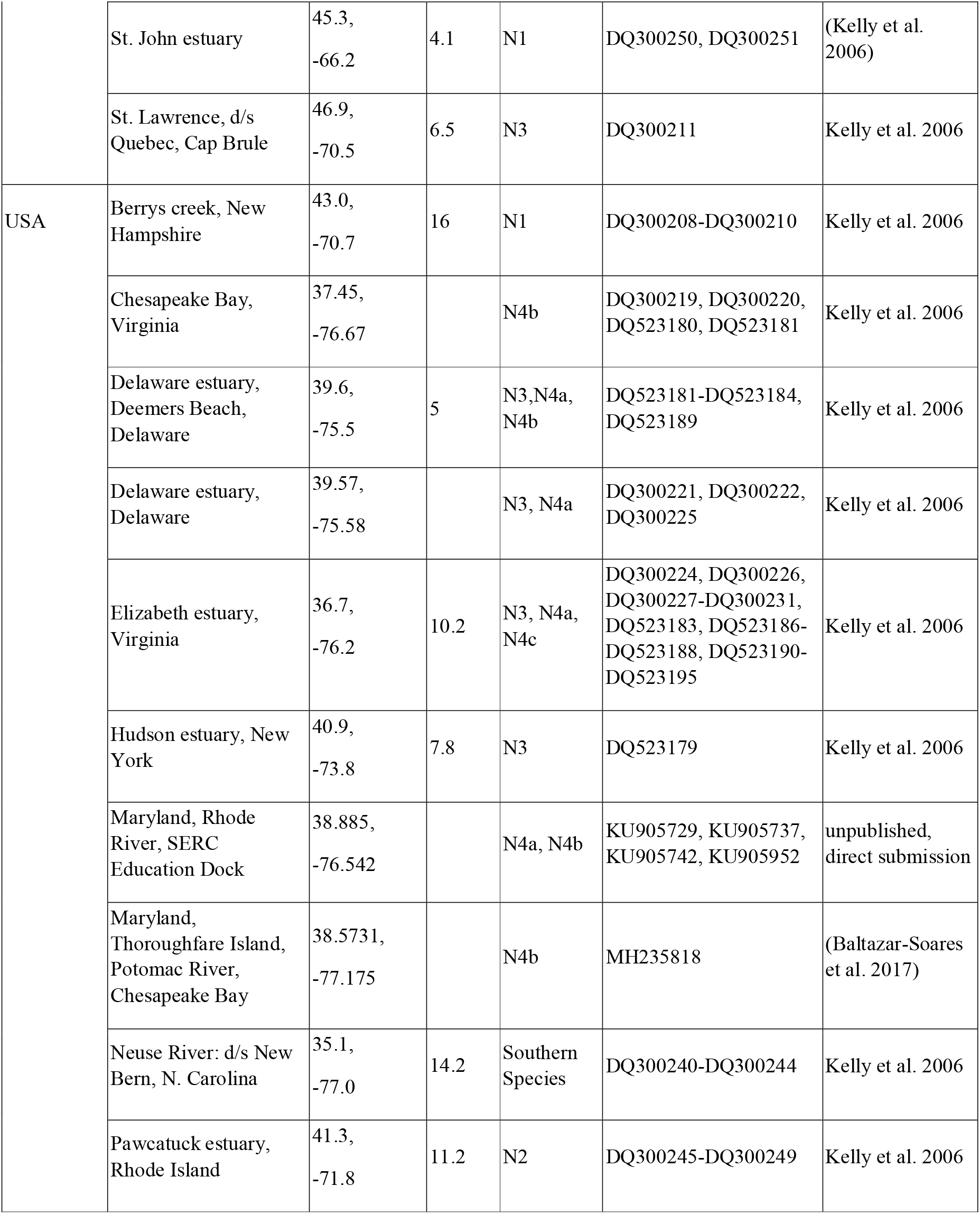
Sample list of specimens used for the phylogenetic tree and the determination of haplotypes. Indicated are the localities, if available the salinity of the sampling site, the identified haplotypes and the accession numbers (F=freshwater sites).

**Supplement Table 2:** Differences of biomass change between control group (without G. tigrinus) and treatment group (with G. tigrinus) with respect to species, Salinity (S) and temperature (T). Furthermore the differences between the groups were statistical analyzed and the p-value calculated with the non-parametrical Mann-Whitney U-Test.

| Species | S<br>[g·kg<sup>-1</sup>] | T<br>[°C] | N | Median<br>Control | Median<br>Treatment | Δ Median<br>(T-C) | p-value<br>(U-Test) |
| --- | --- | --- | --- | --- | --- | --- | --- |
| *Chara aspera* | 3 | 9 | 8 | -0,7 | -11,0 | -10,3 | 0,01* |
| *Chara aspera* | 3 | 15 | 8 | -0,7 | -13,7 | -13,0 | 0,01* |
| *Chara aspera* | 3 | 24 | 8 | -7,3 | -26,0 | -18,7 | 0,03* |
| *Chara aspera* | 3 | 30 | 8 | -15,3 | -11,8 | 3,5 | 0,60 |
| *Chara aspera* | 5 | 9 | 8 | -0,5 | -9,0 | -8,5 | 0,01* |
| *Chara aspera* | 5 | 15 | 8 | -2,9 | -15,2 | -12,2 | 0,01* |
| *Chara aspera* | 5 | 24 | 8 | -2,3 | -20,1 | -17,8 | 0,01* |
| *Chara aspera* | 5 | 30 | 8 | -3,3 | -28,5 | -25,2 | 0,00* |
| *Chara aspera* | 7 | 9 | 8 | 0,2 | -7,6 | -7,8 | 0,12 |
| *Chara aspera* | 7 | 15 | 8 | 0,4 | -18,5 | -18,8 | 0,01* |
| *Chara aspera* | 7 | 24 | 8 | -2,2 | -33,6 | -31,3 | 0,00* |
| *Chara aspera* | 7 | 30 | 8 | -12,9 | -34,9 | -22,0 | 0,03* |
| *Chara aspera* | 13 | 9 | 8 | 0,9 | -5,9 | -6,8 | 0,02* |
| *Chara aspera* | 13 | 15 | 8 | 0,2 | -13,5 | -13,7 | 0,02* |
| *Chara aspera* | 13 | 24 | 8 | -4,2 | -35,3 | -31,1 | 0,01* |
| *Chara aspera* | 13 | 30 | 8 | -14,3 | -34,0 | -19,8 | 0,02* |
| *Chara tomentosa* | 3 | 9 | 8 | -0,7 | -2,6 | -1,9 | 0,17 |
| *Chara tomentosa* | 3 | 15 | 8 | -2,8 | -7,3 | -4,5 | 0,12 |
| *Chara tomentosa* | 3 | 24 | 8 | -2,0 | -17,3 | -15,4 | 0,01* |
| *Chara tomentosa* | 3 | 30 | 8 | -4,7 | -14,5 | -9,7 | 0,04* |
| *Chara tomentosa* | 5 | 9 | 8 | 0,9 | -5,5 | -6,4 | 0,04* |
| *Chara tomentosa* | 5 | 15 | 8 | -0,3 | -9,0 | -8,7 | 0,14 |
| *Chara tomentosa* | 5 | 24 | 8 | -3,6 | -8,2 | -4,6 | 0,03* |
| *Chara tomentosa* | 5 | 30 | 8 | -3,3 | -11,1 | -7,9 | 0,03* |
| *Chara tomentosa* | 7 | 9 | 8 | -1,7 | -6,0 | -4,3 | 0,12 |
| *Chara tomentosa* | 7 | 15 | 8 | 3,2 | -6,5 | -9,7 | 0,01* |
| *Chara tomentosa* | 7 | 24 | 8 | 2,0 | -17,0 | -19,0 | 0,04* |
| *Chara tomentosa* | 7 | 30 | 8 | -7,0 | -9,6 | -2,6 | 0,35 |
| *Chara tomentosa* | 13 | 9 | 8 | -0,9 | -5,7 | -4,8 | 0,00* |
| *Chara tomentosa* | 13 | 15 | 8 | -0,1 | -7,5 | -7,4 | 0,25 |
| *Chara tomentosa* | 13 | 24 | 8 | -1,5 | -6,7 | -5,2 | 0,02* |
| *Chara tomentosa* | 13 | 30 | 8 | -4,0 | -13,7 | -9,7 | 0,02* |
| *Chara sp.* | 3 | 9 | 16 | -0,7 | -7,5 | -6,8 | 0,01* |
| *Chara sp.* | 3 | 15 | 16 | -1,7 | -8,6 | -6,9 | 0,00* |
| *Chara sp.* | 3 | 24 | 16 | -3,1 | -17,7 | -14,6 | 0,00* |
| *Chara sp.* | 3 | 30 | 16 | -6,1 | -14,0 | -7,8 | 0,03* |
| *Chara sp.* | 5 | 9 | 16 | 0,2 | -8,3 | -8,4 | 0,00* |
| *Chara sp.* | 5 | 15 | 16 | -1,7 | -10,4 | -8,7 | 0,00* |
| *Chara sp.* | 5 | 24 | 16 | -2,6 | -14,9 | -12,4 | 0,00* |
| *Chara sp.* | 5 | 30 | 16 | -3,3 | -15,5 | -12,2 | 0,00* |
| *Chara sp.* | 7 | 9 | 16 | -1,1 | -6,2 | -5,1 | 0,03* |
| *Chara sp.* | 7 | 15 | 16 | 1,8 | -8,3 | -10,0 | 0,00* |
| *Chara sp.* | 7 | 24 | 16 | 1,2 | -22,9 | -24,2 | 0,00* |
| *Chara sp.* | 7 | 30 | 16 | -10,7 | -20,4 | -9,6 | 0,04* |
| *Chara sp.* | 13 | 9 | 16 | 0,6 | -5,9 | -6,6 | 0,00* |
| *Chara sp.* | 13 | 15 | 16 | 0,0 | -9,9 | -9,9 | 0,01* |
| *Chara sp.* | 13 | 24 | 16 | -2,3 | -29,1 | -26,9 | 0,00* |
| *Chara sp.* | 13 | 30 | 16 | -8,1 | -20,0 | -11,8 | 0,01* |

**Figure S1.**
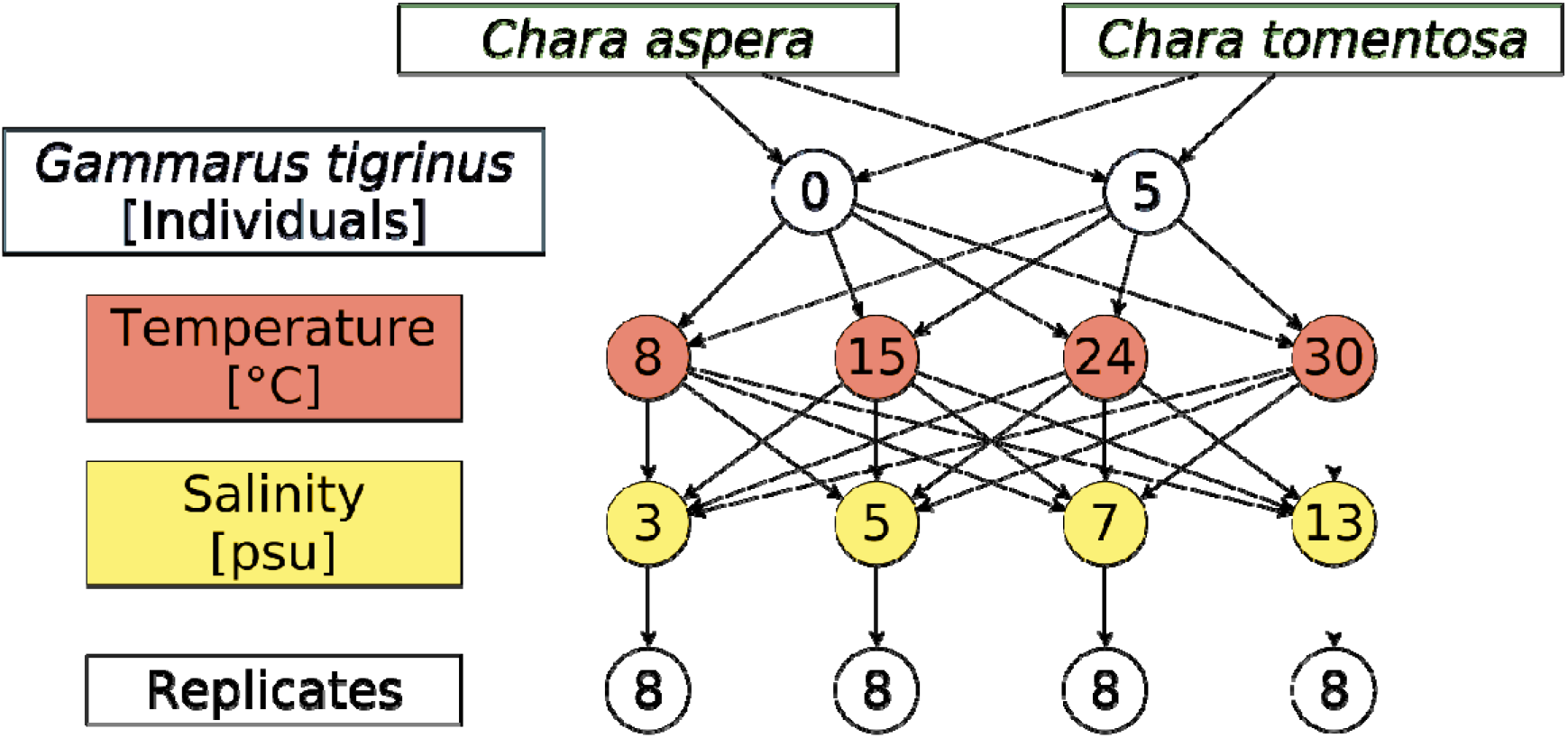
Flow scheme of the full factorial experimental design for temperature, and salinity with *Chara aspera* and *C. tomentosa*, with and without *Gammarus tigrinus*.

**Figure S2.**
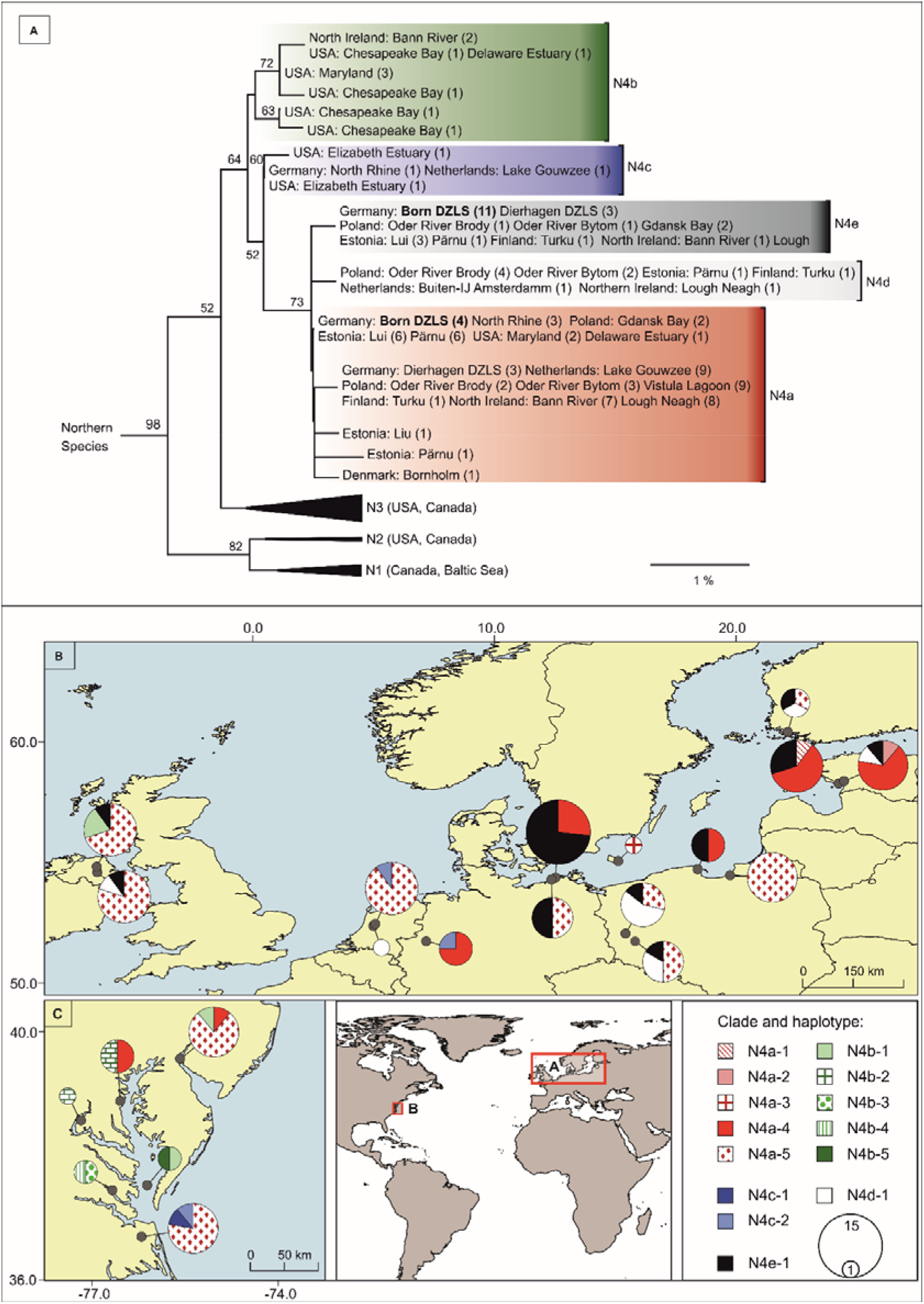
(A) Phylogeny of *Gammarus tigrinus* COI haplotypes inferred by using the Maximum Likelihood method (Tamura-Nei model +I+Γ) with bootstrap values above the branches (1000 replicates). The clade numbers are identical to Kelly et al. (2006) and the coloured boxes indicate haplotypes of clade N4. Haplotypes N4a and N4e comprised the individuals from the DZLS. (B & C) Distribution of *Gammarus tigrinus* haplotypes of clade N4 identified by COI network analysis. Pie charts indicated haplotype frequencies in European (B) and North American (C) populations. More information on haplotypes can be found in Supplement Table S1.

**Figure S3.**
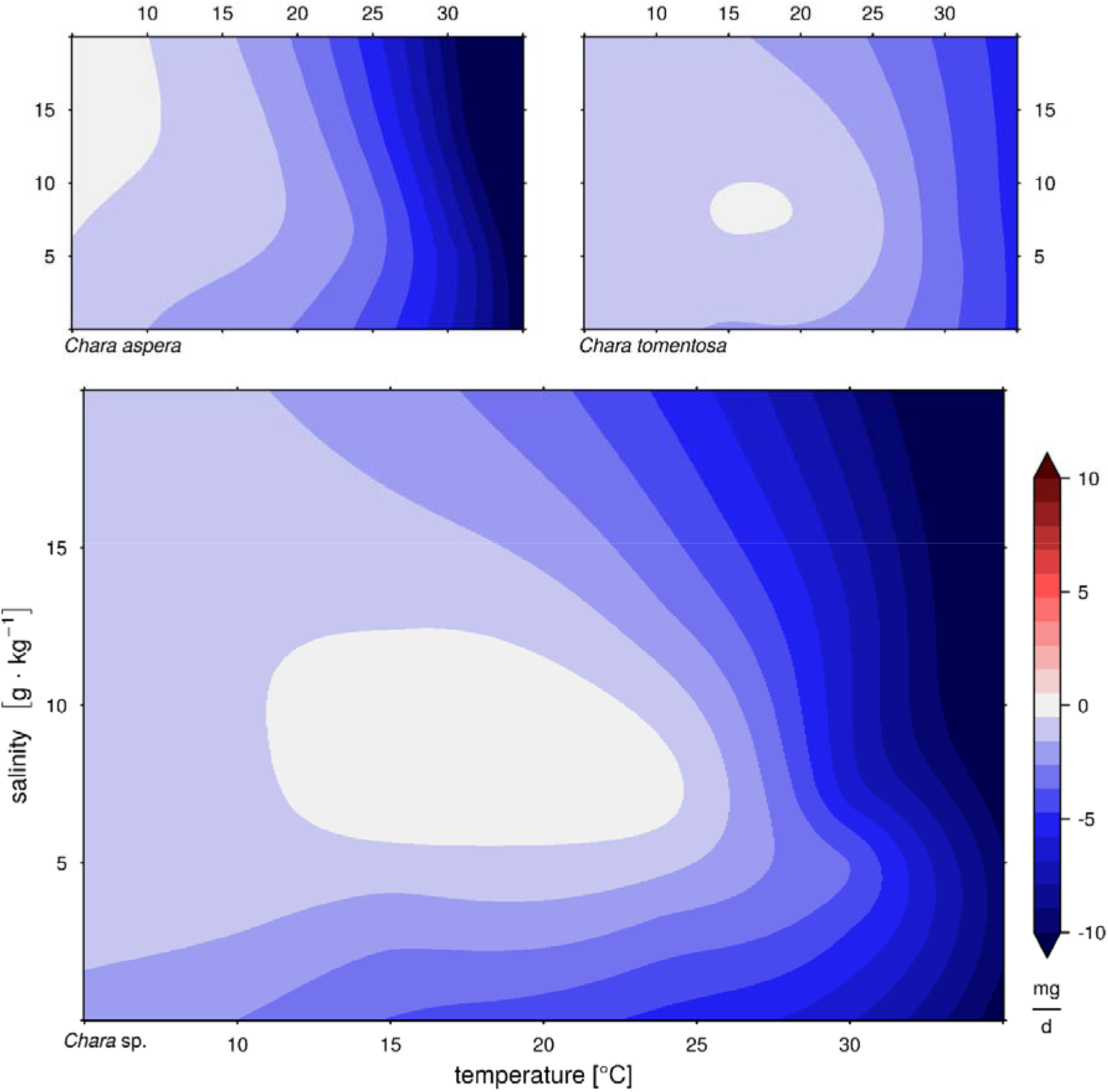
Generalized additive model for change in biomass of *Chara aspera* (top left), *Chara tomentosa* (top right) and for both Chara species (bottom) in respect to different salinity and temperature levels.

**Figure S4.**
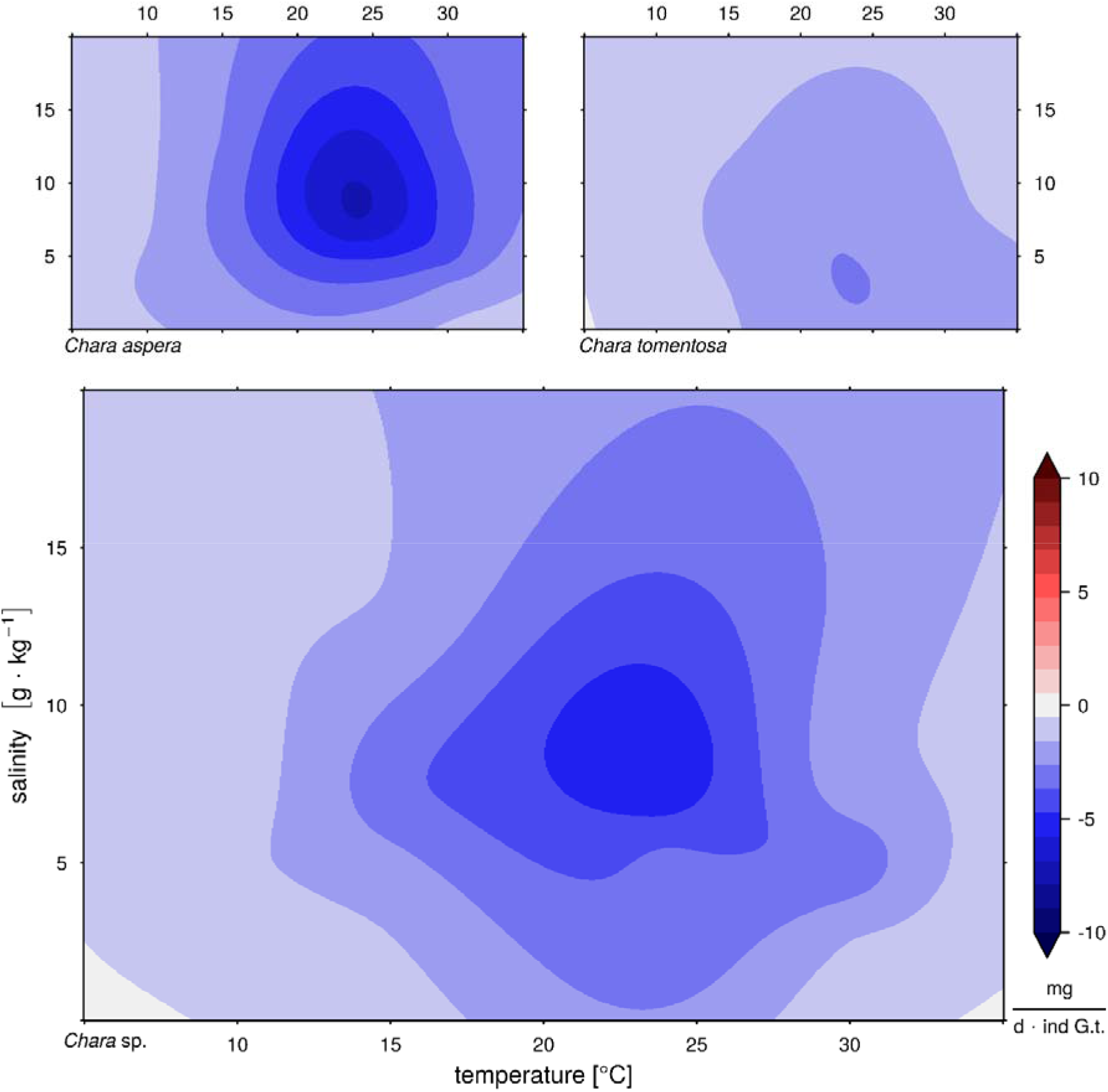
Generalized additive model for loss of biomass caused by growth corrected grazing of an individual of *Gammarus tigrinus* (G.t.) on *Chara aspera* (top left), *Chara tomentosa* (top right) and on both Chara species (bottom) in respect to different salinity and temperature levels.

**Figure S5.**
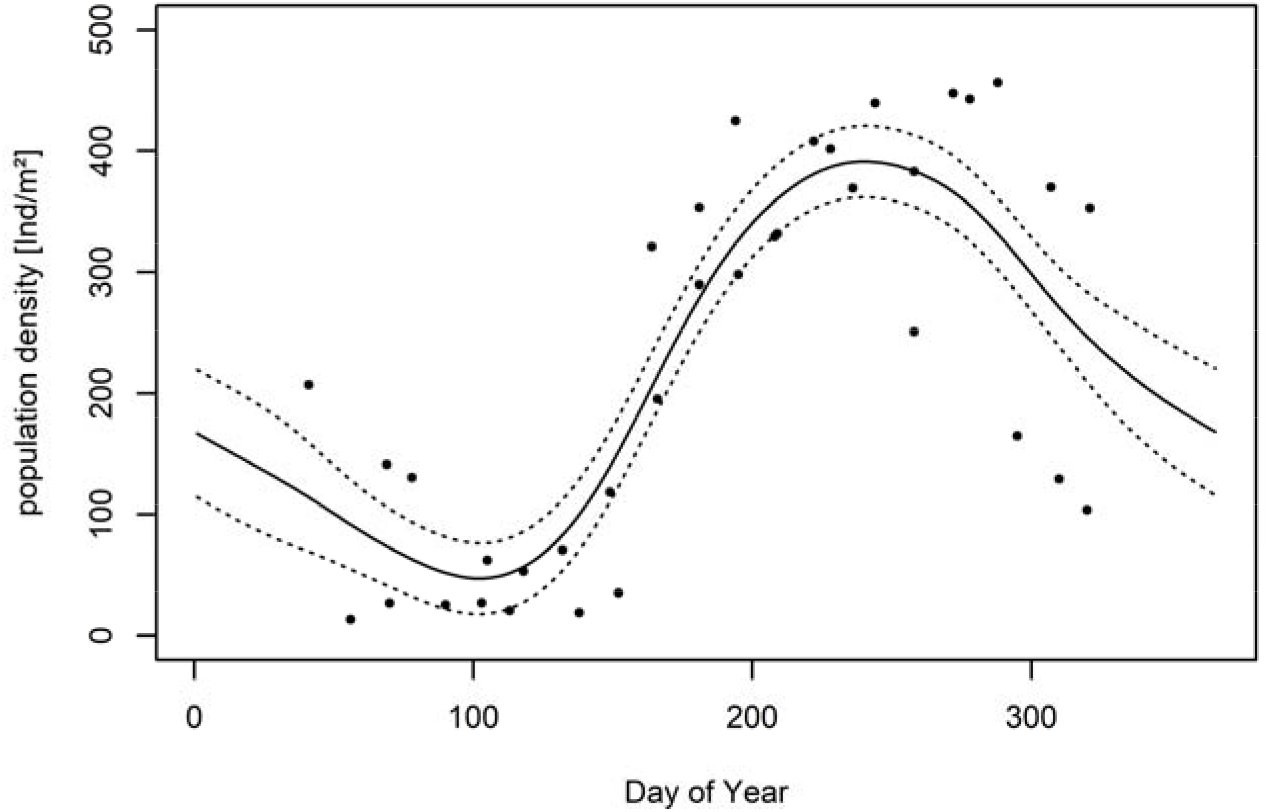
Model of seasonal pattern of *Gammarus tigrinus* calculated with population densities from Chambers (1977) and modified to have a maximum of 400 individuals per square meter based on values of Zettler (1995) and of M. Paar (personal communication).

## Elemental composition

There were no significant differences for the C:N and C:P ratios along the depth distribution, or between stations (see pooled values Figure S6). Charophytes showed either a weak seasonal trend in elemental composition (C:N), or the ratio decreased (C:P). Lowest C:P ratios were found from July to October. Contrary, co-occurring tracheophytes increased their C:N ratio during summer, as well as their C:P ratios. These findings indicate that charophytes had probably a higher protein content per biomass compared to tracheophytes.

**Figure S6.**
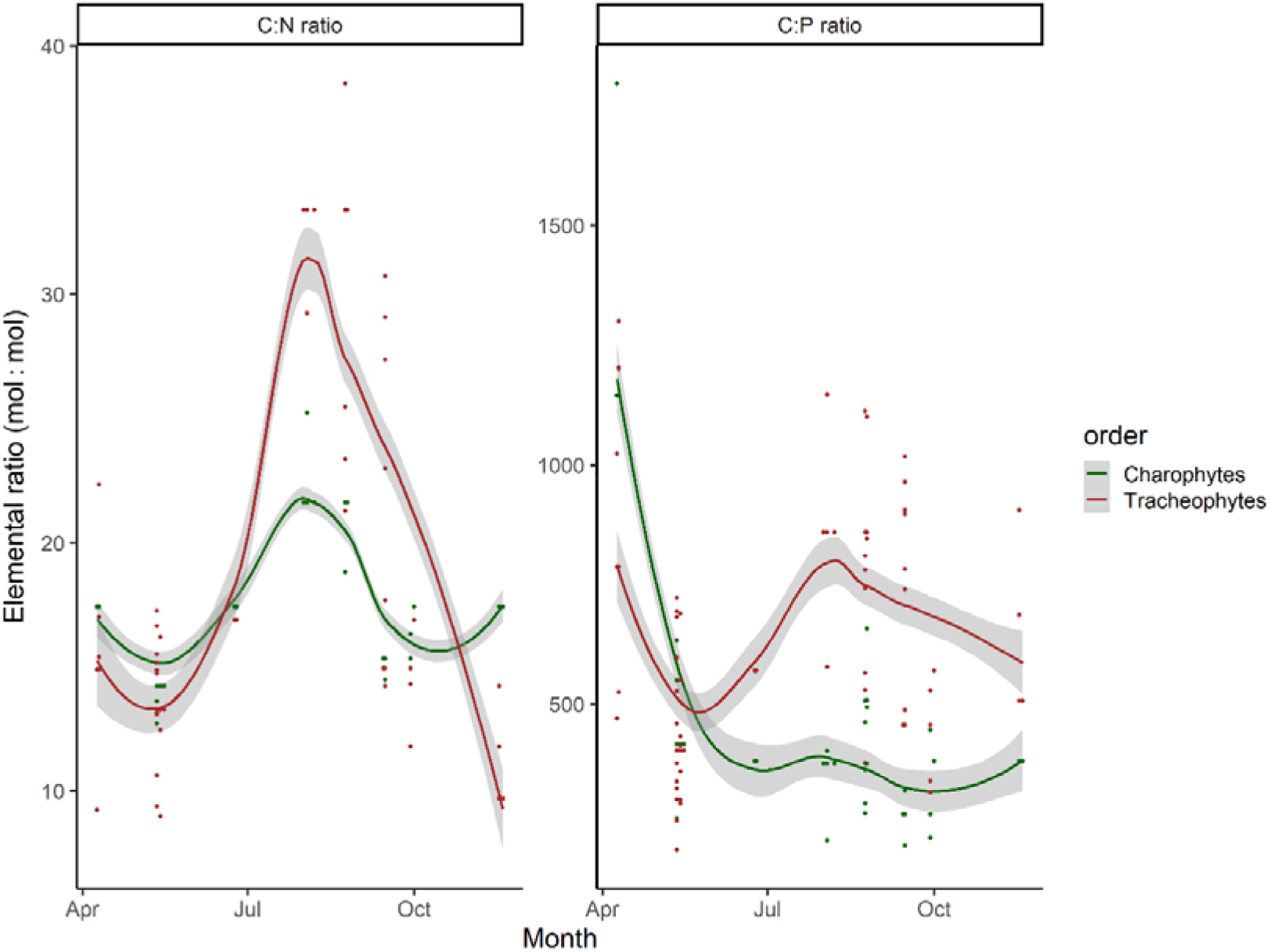
Elemental ratio of Carbon to Nitrogen (C:N) and Carbon to Phosphorus (C:P) of macrophyte biomass (separated by order) at two stations of the Darss-Zingst lagoon system across all sampling stations (Bodstedter Bodden and Grabow) and sampling depths in 2014. Please note the different scaling. Points represent measured elemental ratios, lines a local regression (LOESS – locally estimated scatter smoothing), and the ribbons the 95% confidence intervals (Wickham 2016).

## References

Auderset Joye, D., and A. Rey-Boissezon. 2015. Will charophyte species increase or decrease their distribution in a changing climate? Aquatic Botany 120:73–83.

Auderset Joye, D., and A. Schwarzer. 2012. Red List Characeae. Threatened Species in Switzerland. Status 2010.

Baltazar-Soares, M., F. Paiva, Y. Chen, A. Zhan, and E. Briski. 2017. Diversity and distribution of genetic variation in gammarids: Comparing patterns between invasive and non-invasive species. Ecology and Evolution 7:7687–7698.

Bandelt, H. J., P. Forster, and A. Rohl. 1999. Median-joining networks for inferring intraspecific phylogenies. Molecular Biology and Evolution 16:37–48.

Berezina, N. A. 2007. Expansion of the North American amphipod Gammarus tigrinus Sexton, 1939 to the Neva Estuary (easternmost Baltic Sea). Oceanologia 49:129–135.

Berthold, M., U. Karsten, M. von Weber, A. Bachor, and R. Schumann. 2018a. Phytoplankton can bypass nutrient reductions in eutrophic coastal water bodies. Ambio 47 (1):146–158.

Berthold, M., S. Karstens, U. Buczko, and R. Schumann. 2018b. Potential export of soluble reactive phosphorus from a coastal wetland in a cold-temperate lagoon system: Buffer capacities of macrophytes and impact on phytoplankton. Science of the Total Environment 616–617:46–54.

Berthold, M., D. Zimmer, V. Reiff, and R. Schumann. 2018c. Phosphorus Contents Re-visited After 40 Years in Muddy and Sandy Sediments of a Temperate Lagoon System. Frontiers in Marine Science 5:1–14.

Berthold, M., D. Zimmer, and R. Schumann. 2015. A simplified method for total phosphorus digestion with potassium persulphate at sub-boiling temperatures in different environmental samples. Rostocker Meeresbiologische Beiträge 25:7–25.

Björkman, S. O. 1947. On the distribution of Chara tomentosa L. round the Baltic and some remarks on its specific epithet. Botaniska Notiser 1947:157–170.

Blazencic, J., B. Stevanovic, Z. Blazencic, and V. Stevanovic. 2005. Red Data List of Charophytes in the Balkans. Biodiversity & Conservation 15:3445–3457.

Blindow, I. 2000. Distribution of charophytes along the Swedish coastin relation to salinity and eutrophication. International Review of Hydrobiology 85:707–717.

Blindow, I., S. Dahlke, A. Dewart, S. Flügge, M. Hendreschke, A. Kerkow, and J. Meyer. 2016. Long-term and interannual changes and viable diaspore reservoir of submerged macrophytes in a shallow brackish water bay of the southern Baltic Sea - influence of eutrophication and climate. Hydrobiologia:1–16.

Blindow, I., J. Dietrich, N. Möllmann, and H. Schubert. 2003. Growth, photosynthesis and fertility of Chara aspera under different light and salinity conditions. Aquatic Botany 76:213–234.

Blindow, I., N. Möllmann, M. G. Boegle, and M. Schütte. 2009. Reproductive isolation in Chara aspera populations. Aquatic Botany 91:224–230.

Blindow, I., and M. Schütte. 2007. Elongation and mat formation of Chara aspera under different light and salinity conditions. Hydrobiologia 584:69–76.

Buchsbaum, R., I. Valiela, T. Swain, M. Dzierzeski, and S. Allen. 1991. Available and refractory nitrogen in detritus of coastal vascular plants and macroalgae. Marine Ecology Progress Series 72:131–143.

Chambers, M. R. 1977. The population ecology of Gammarus tigrinus (sexton) in the reed beds of the Tjeukemeer. Hydrobiologia 53:155–164.

Choudhury, M. I., P. Urrutia-Cordero, H. Zhang, M. K. Ekvall, L. R. Medeiros, and L.-A. Hansson. 2019. Charophytes collapse beyond a critical warming and brownification threshold in shallow lake systems. Science of The Total Environment 661:148–154.

Daunys, D., and M. L. Zettler. 2006. Invasion of the North Amrican Amphipod (Gammarus tigrinus Sexton, 1939) into the Curonian Lagoon, South-Eastern Baltic Sea. Acta Zoologica Lituanica 16:20–26.

Dorgelo, J. 1973. Comparative ecophysiology of gammarids (crustacea: amphipoda) from marine, brackish and fresh-water habitats exposed to the influence of salinity- temperature combinations. III. Oxygen uptake. Netherlands Journal of Sea Research 7:253–266.

Dorgelo, J. 1974. Comparative ecophysiology of gammarids (Crustacea: Amphipoda) from marine, brackish and fresh-water habitats, exposed to the influence of salinity- temperature combinations - I. Effect on survival. Hydrobiological Bulletin 8:90–108.

Felten, V., G. Tixier, F. Guérold, V. De Crespin De Billy, O. Dangles, F. Guerold, De Crespin V. De Billy, O. Dangles, F. Guérold, V. De Crespin De Billy, and O. Dangles. 2008. Quantification of diet variability in a stream amphipod: Implications for ecosystem functioning. Fundamental and Applied Limnology / Archiv für Hydrobiologie 170:303– 313.

Folmer, O., M. Black, W. Hoeh, R. Lutz, and R. Vrijenhoek. 1994. DNA primers for amplification of mitochondrial cytochrome c oxidase subunit I from diverse metazoan invertebrates. Molecular marine biology and biotechnology 3:294–299.

Fox, J., and S. Weisberg. 2019. An {R} Companion to Applied Regression. Third. Sage, Thousand Oaks {CA}.

Gärdenfors, U. 2010. 2010 Rödlistade arten I Sverige (The 2010 Red List of Swedish Species). ArtDatabanken. Uppsala.

Gessner, F. 1957. Meer und Strand. Page (F. Gessner, Ed.). Second edition. VEB Deutscher Verlag der Wissenschaften, Berlin.

Grabner, D. S., A. M. Weigand, F. Leese, C. Winking, D. Hering, R. Tollrian, and B. Sures. 2015. Invaders, natives and their enemies: distribution patterns of amphipods and their microsporidian parasites in the Ruhr Metropolis, Germany. Parasites and Vectors 8:1– 15.

Hasslow, O. J. 1931. Sveriges characéer. Botaniska Notiser:63–136.

HELCOM. 2010. Ecosystem Health of the Baltic sea 2003-2007: HELCOM Initial Holistic Assessment. Baltic Sea Environment Proceedings 122:68.

HELCOM. 2018. Sources and pathways of nutrients to the Baltic Sea. Baltic Sea Environmental Proceedings 153:48.

Herkül, K., V. Lauringson, and J. Kotta. 2016. Specialization among amphipods: The invasive Gammarus tigrinus has narrower niche space compared to native gammarids. Ecosphere 7:1–16.

Janssen, F., T. Neumann, and M. Schmidt. 2004. Inter-annual variability in cyanobacteria blooms in the Baltic Sea controlled by wintertime hydrographic conditions. Marine Ecology Progress Series 275:59–68.

Karstens, S., U. Buczko, and S. Glatzel. 2015. Phosphorus storage and mobilization in coastal Phragmites wetlands: Influence of local-scale hydrodynamics. Estuarine, Coastal and Shelf Science 164:124–133.

Karstens, S., U. Buczko, G. Jurasinski, R. Peticzka, and S. Glatzel. 2016. Impact of adjacent land use on coastal wetland sediments. Science of the Total Environment 550:337–348.

Kelley, A. L. 2014. The role thermal physiology plays in species invasion. Conservation Physiology 2:1–14.

Kelly, D. W., J. T. A. Dick, and W. I. Montgomery. 2002. The functional role of Gammarus (Crustacea, Amphipoda): Shredders, predators, or both? Hydrobiologia 485:199–203.

Kelly, D. W., J. R. Muirhead, D. D. Heath, and H. J. Macisaac. 2006. Contrasting patterns in genetic diversity following multiple invasions of fresh and brackish waters. Molecular Ecology 15:3641–3653.

Koop, J. H. E., and M. K. Grieshaber. 2000. The role of ion regulation in the control of the distribution of Gammarus tigrinus (Sexton) in salt-polluted rivers. Journal of Comparative Physiology - B Biochemical, Systemic, and Environmental Physiology 170:75–83.

Körner, S., and T. Dugdale. 2003. Is roach herbivory preventing re-colonization of submerged macrophytes in a shallow lake? Hydrobiologia 506–509:497–501.

Korpinen, S., and M. Westerbom. 2009. Microhabitat segregation of the amphipod genus Gammarus (Crustacea: Amphipoda) in the Northern Baltic Sea. Marine Biology 157:361–370.

Kotta, J., H. Orav-Kotta, K. Herkül, and I. Kotta. 2011. Habitat choice of the invasive *Gammarus tigrinus* and the native *Gammarus salinus* indicates weak interspecific competition. Boreal Environment Research 16:64–72.

Kraufvelin, P., S. Salovius, H. Christie, F. E. Moy, R. Karez, and M. F. Pedersen. 2006. Eutrophication-induced changes in benthic algae affect the behaviour and fitness of the marine amphipod Gammarus locusta. Aquatic Botany 84:199–209.

Kufel, L., and I. Kufel. 2002. Chara beds acting as nutrient sinks in shallow lakes—a review. Aquatic Botany 72:249–260.

Kumar, S., G. Stecher, M. Li, C. Knyaz, and K. Tamura. 2018. MEGA X: Molecular evolutionary genetics analysis across computing platforms. Molecular Biology and Evolution 35:1547–1549.

Kuprijanov, I., J. Kotta, V. Lauringson, and K. Herkül. 2015. Trophic interactions between native and alien palaemonid prawns and an alien gammarid in a brackish water ecosystem. Proceedings of the Estonian Academy of Sciences 64:518–524.

Leigh, J. W., and D. Bryant. 2015. POPART: Full-feature software for haplotype network construction. Methods in Ecology and Evolution 6:1110–1116.

Lenz, M., B. A. P. P. da Gama, N. V. Gerner, J. Gobin, F. Gröner, A. Harry, S. R. Jenkins, P. Kraufvelin, C. Mummelthei, J. Sareyka, E. A. Xavier, and M. Wahl. 2011. Non-native marine invertebrates are more tolerant towards environmental stress than taxonomically related native species: results from a globally replicated study. Environmental Research 111:943–52.

van der Linden, P., and J. F. B. Mitchell. (n.d.). ENSEMBLES: Climate Change and its Impacts: Summary of research and results from the ENSEMBLES project. FitzRoy Road, Exeter EX1 3PB, UK.

Lindner, A. 1972. Soziologisch-ökologische Untersuchungen an der submersen Vegetation in der Boddenkette südlich des Darß und Zingst. University of Rostock.

Melzer, A. 1999. Aquatic macrophytes as tools for lake management. Hydrobiologia 395/396:181–190.

Neumann, T., and R. Friedland. 2011. Climate Change Impacts on the Baltic Sea. Pages 23– 32 in G. Schernewski, J. Hofstede, and T. Neumann, editors. Global Change and Baltic Coastal Zones. Springer.

Normant, M., M. Feike, A. Szaniawska, and G. Graf. 2007. Adaptation of Gammarus tigrinus Sexton, 1939 to new environments-Some metabolic investigations. Thermochimica Acta 458:107–111.

Orav-Kotta, H., J. Kotta, K. Herkül, I. Kotta, and T. Paalme. 2009. Seasonal variability in the grazing potential of the invasive amphipod Gammarus tigrinus and the native amphipod Gammarus salinus (Amphipoda: Crustacea) in the northern Baltic Sea. Biological Invasions 11:597–608.

Östman, Ö., J. Eklöf, B. K. Eriksson, J. Olsson, P. O. Moksnes, and U. Bergström. 2016. Top- down control as important as nutrient enrichment for eutrophication effects in North Atlantic coastal ecosystems. Journal of Applied Ecology 53:1138–1147.

Paar, M., M. Berthold, R. Schumann, S. Dahlke, and I. Blindow. 2021. Seasonal Variation in Biomass and Production of the Macrophytobenthos in two Lagoons in the Southern Baltic Sea. Frontiers in Earth Science 8:1–16.

Paiva, F., A. Barco, Y. Chen, A. Mirzajani, F. T. Chan, V. Lauringson, M. Baltazar-Soares, A. Zhan, S. A. Bailey, J. Javidpour, and E. Briski. 2018. Is salinity an obstacle for biological invasions? Global Change Biology 24:2708–2720.

Pellan, L., V. Médoc, D. Renault, T. Spataro, and C. Piscart. 2016. Feeding choice and predation pressure of two invasive gammarids, Gammarus tigrinus and Dikerogammarus villosus, under increasing temperature. Hydrobiologia 781:43–54.

Piepho, M. 2017. Assessing maximum depth distribution, vegetated area, and production of submerged macrophytes in shallow, turbid coastal lagoons of the southern Baltic Sea. Hydrobiologia 794:303–316.

Pitkänen, H., M. Peuraniemi, M. Westerbom, M. Kilpi, and M. Von Numers. 2013. Long- term changes in distribution and frequency of aquatic vascular plants and charophytes in an estuary in the Baltic Sea. Annales Botanici Fennici 50:1–54.

Pohl, P., M. K. Ohlhase, S. K. Rautwurst, and K. Laus-Kinnerk Baasch. 1987. An inexpensive inorganic medium for the mass cultivation of freshwater microalgae. Phytochemistry 26:1657–1659.

Puche, E., S. Sánchez-Carrillo, M. Álvarez-Cobelas, A. Pukacz, M. A. Rodrigo, and C. Rojo. 2018. Effects of overabundant nitrate and warmer temperatures on charophytes: The roles of plasticity and local adaptation. Aquatic Botany 146:15–22.

Puntila, R. 2016. Trophic Interactions and Impacts of Non-indigenous Species in Baltic Sea Coastal Ecosystems. University of Helsinki.

R Core Team. 2019. R: A Language and Environment for Statistical Computing. Vienna, Austria.

Radulovici, A. E., B. Sainte-Marie, and F. Dufresne. 2009. DNA barcoding of marine crustaceans from the Estuary and Gulf of St Lawrence: A regional-scale approach. Molecular Ecology Resources 9:181–187.

Reusch, T. B. H., J. Dierking, H. C. Andersson, E. Bonsdorff, J. Carstensen, M. Casini, M. Czajkowski, B. Hasler, K. Hinsby, K. Hyytiäinen, K. Johannesson, S. Jomaa, V. Jormalainen, H. Kuosa, S. Kurland, L. Laikre, B. R. MacKenzie, P. Margonski, F. Melzner, D. Oesterwind, H. Ojaveer, J. C. Refsgaard, A. Sandström, G. Schwarz, K. Tonderski, M. Winder, and M. Zandersen. 2018. The Baltic Sea as a time machine for the future coastal ocean. Science Advances 4:eaar8195.

Rojo, C., M. Carramiñana, D. Cócera, G. P. Roberts, E. Puche, S. Calero, and M. A. Rodrigo. 2017. Different responses of coexisting Chara species to foreseeable Mediterranean temperature and salinity increases. Aquatic Botany in press.

Sanudo-Wilhelmy, S. A., A. Tovar-Sanchez, F. Fu, D. G. Capone, E. J. Carpenter, and D. A. Hutchins. 2004. The impact of surface-adsorbed phosphorus on phytoplankton Redfield stoichiometry. Nature 432:897–901.

Schiewer, U. 2007. Darß-Zingst Boddens, Northern Rügener Boddens and Schlei. Pages 35– 86 in U. Schiewer, M. M. Caldwell, G. Heldmaier, R. B. Jackson, O. L. Lange, H. A. Mooney, E.-D. Schulze, and U. Sommer, editors. Ecology of Baltic coastal waters. First edition. Springer, Berlin, Heidelberg.

Schubert, H., and I. Blindow, editors. 2003. Charophytes of the Baltic Sea. Gantner, Ruggel, Königstein and Germany.

Schubert, H., I. Blindow, N. C. Bueno, M. T. Casanova, M. Pełechaty, and A. Pukacz. 2018. Ecology of charophytes – permanent pioneers and ecosystem engineers. Perspectives in Phycology 5:61–74.

Schumann, R., H. Baudler, Ä. Glass, K. Dümcke, and U. Karsten. 2006. Long-term observation on salinity dynamics in a tideless shallow coastal lagoon of the Southern Baltic Sea coast and their biological relevance. Journal of Marine Systems 60:330–344.

Stewart, N. F., and J. M. Church. 1992. Red data books of Britain and Ireland-- Stoneworts. {Joint Nature Conservation Committee} and {Office of Public Works} and Plantlife, Peterborough.

Torn, K., G. Martin, and R. Munsterhjelm. 2003. Chara tomentosa L. 1753. Pages 131–141 in H. Schubert and I. Blindow, editors. Charophytes of the Baltic Sea. Gantner, Ruggel, Königstein, Germany.

Torn, K., G. Martin, and T. Paalme. 2006. Seasonal changes in biomass, elongation growth and primary production rate of Chara tomentosa in the NE Baltic Sea. Annales Botanici Fennici 43:276–283.

Volkmann, C. 2016. The impact of organic material for macrophytes in coastal waters of the Baltic Sea. University of Rostock.

Wickham, H. 2016. ggplot2: Elegant Graphics for Data Analysis. Springer-Verlag New York.

Wood, S. N. 2011. Fast stable restricted maximum likelihood and marginal likelihood estimation of semiparametric generalized linear models. Journal of the Royal Statistical Society (B) 73:3–36.

Wüstenberg, A., Y. Pörs, and R. Ehwald. 2011. Culturing of stoneworts and submersed angiosperms with phosphate uptake exclusively from an artificial sediment. Freshwater Biology 56:1531–1539.

Zettler, M. L. 1995. Erstnachweis von Gammarus tigrinus Sexton, 1939 (Crustacea: Amphipoda) in der Darss-Zingster-Boddenkette und seine derzeitige Verbreitung an der deutschen Ostsekueste. Arch. Freunde Naturg. Mecklb. XXXIV:137–141.

Zettler, M. L. 2001. Some malacostracan crustacean assemblages in the southern and western Baltic Sea. Rostock. Meersebiolog. Beitr. 9:127–143.

Zettler, M. L., and A. Zettler. 2017. Marine and freshwater Amphipoda from the Baltic Sea and adjacent territories. 2. Auflage. Conchbooks, Hackenheim.

